# Chloroplastic ascorbate acts as a regulatory hub in plant metabolism regardless of oxidative stress

**DOI:** 10.1101/2024.03.14.585081

**Authors:** Dávid Tóth, Roland Tengölics, Fayezeh Aarabi, Anna Karlsson, André Vidal-Meireles, László Kovács, Soujanya Kuntam, Tímea Körmöczi, Alisdair R. Fernie, Elton P. Hudson, Balázs Papp, Szilvia Z. Tóth

**Affiliations:** Laboratory for Molecular Photobioenergetics, Institute of Plant Biology, HUN-REN Biological Research Centre, Szeged, Temesvári krt 62, H-6726 Szeged, Hungary; Doctoral School of Biology, University of Szeged, Közép fasor 52, H-6722 Szeged, Hungary; HCEMM-BRC Metabolic Systems Biology Lab; Temesvári krt 62, H-6726, Szeged, Hungary; Synthetic and Systems Biology Unit, Institute of Biochemistry, HUN-REN Biological Research Centre, Szeged, Temesvári krt 62, H-6726 Szeged, Hungary; Metabolomics Lab, Core facilities, HUN-REN Biological Research Centre, Temesvári krt 62, H-6726 Szeged, Hungary; Max-Planck-Institute of Molecular Plant Physiology, Am Mühlenberg 1, D-14476 Potsdam-Golm, Germany; Science for Life Laboratory, School of Engineering Science in Chemistry, Biotechnology and Health, KTH Royal Institute of Technology, P-Box 1031, 171 21, Solna, Sweden; National Laboratory for Health Security, HUN-REN Biological Research Centre, Szeged, Temesvári krt 62, H-6726 Szeged Hungary

**Keywords:** Ascorbate, GDP-L-galactose phosphorylase, PHT4;4 transporter, Proteome Integral Solubility Alteration assay, untargeted metabolomics

## Abstract

Ascorbate is a major plant metabolite that plays crucial roles in various processes, from reactive oxygen scavenging to epigenetic regulation. However, to what extent and how ascorbate modulates metabolism is largely unknown. To address this, we investigated the consequences of chloroplastic and total cellular ascorbate-deficiencies by studying chloroplastic ascorbate-transporter *pht4;4* mutant lines, and the ascorbate-deficient *vtc2-4* mutant of *Arabidopsis thaliana*. Under regular growth conditions, both ascorbate-deficiencies caused minor alterations in photosynthesis, with no apparent signs of oxidative damage. In contrast, metabolomics analysis revealed a global and largely overlapping metabolome rewiring in both ascorbate-deficiencies, suggesting that chloroplastic ascorbate modulates plant metabolism. We observed significant alterations in amino acid metabolism, particularly in arginine metabolism, activation of nucleotide salvage pathways, and changes in secondary metabolism. In addition, proteome-wide analysis of thermostability revealed that ascorbate may interact with enzymes involved in arginine metabolism, the Calvin-Benson cycle, and several photosynthetic electron transport components. Overall, our results suggest that, independently of oxidative stress, chloroplastic ascorbate interconnects and coordinates diverse metabolic pathways in vascular plants and thus acts as a regulatory hub.

## Introduction

Ascorbate (Asc) is a multifunctional metabolite essential for a range of cellular processes in plants, including cell division, expansion, cell wall hydroxylation, signal transduction, programmed cell death, biosynthesis of hormones, iron uptake, epigenetic regulation as a cofactor for DNA and histone demethylases in the nucleus, and reactive oxygen species (ROS) scavenging (reviewed by Noctor et al., 2018; Smirnoff 2018; Tóth et al., 2023). It also acts as a key factor in integrating the interaction of ethylene and abscisic acid and in regulating ROS levels (Yu et al., 2019).

Asc controls stomatal movement in photosynthetic tissues (Chen and Gallie 2004). In the Mehler peroxidase reaction Asc contributes to the detoxification of H_2_O_2_ and the regulation of the redox state of the photosynthetic apparatus (Asada et al., 2006). Within the chloroplast, Asc plays multiple roles (Tóth, 2023). It may act as an alternative electron donor to photosystem II (PSII) and contribute to the plant’s ability to tolerate heat and light stress (Mano et al., 2004; Tóth et al., 2009; Tóth et al., 2011). Asc may also donate electrons to PSII and photosystem I (PSI) in bundle sheath cells with a very low oxygen-evolving capacity (Ivanov et al., 2001; Ivanov et al., 2007). In vascular plants, Asc acts as a cofactor of violaxanthin de-epoxidase, thereby playing an essential role in the process of non-photochemical quenching (NPQ) allowing the dissipation of excess energy as heat (Bratt et al., 1995; Müller-Moulé et al., 2002; Saga et al., 2010; Hallin et al., 2016). On the other hand, Asc-deficiency does not limit NPQ in *Chlamydomonas reinhardtii*, as it is not required as a co-factor for algal-type violaxanthin de-epoxidase (Li et al., 2016; Vidal-Meireles et al., 2020). We also showed that Asc also contributes to dark-induced senescence by inactivating the oxygen-evolving complex (Podmaniczki et al., 2021).

To fulfill the multiple physiological roles of Asc, vascular plants maintain Asc concentrations at high levels of approximately 20 to 30 mM (Zechmann et al., 2011; Zechmann, 2018). Under normal growth conditions, Asc is seemingly in excess, since an 80% reduction of Asc content causes no changes in the phenotype and fails to induce oxidative stress (Müller-Moulé et al., 2003; Müller-Moulé 2004; Lim et al., 2016). On the other hand, under severe environmental stress conditions, Asc may become limited, as shown by an increased oxidative stress tolerance of plants with enhanced Asc regeneration (Wang et al., 2010).

The major route of Asc biosynthesis in plants is the Smirnoff-Wheeler pathway (Wheeler et al., 1998, Smirnoff and Wheeler, 2024), in which *VTC2*, encoding GDP-L-galactose phosphorylase (AT4G26850 in *Arabidopsis thaliana*), plays a central regulatory role (Bulley and Laing 2016). The *vtc2* gene has a homolog, *vtc5*, which provides a minor contribution to Asc biosynthesis (Linster et al., 2008). Other mammalian-type biosynthesis pathways have also been suggested in plants (Wolucka and Montagu, 2003; Agius et al., 2003; Lorence et al., 2004), but the seedling lethality of *vtc2/vtc5* double *A. thaliana* mutants and recent metabolite analysis strongly suggest that their contribution to Asc biosynthesis is negligible or non-existent in *A. thaliana* leaves (Dowdle et al., 2007; Lim et al., 2016; Kavkova et al., 2019; Smirnoff and Wheeler, 2024).

Most of the enzymatic reactions of Asc biosynthesis occur in the cytosol, except the terminal step, catalyzed by L-galactono-1,4-lactone dehydrogenase, which is localized on the inner mitochondrial membrane. From the mitochondria, Asc is transported to the other cell compartments, necessitating specific transporters, since neither Asc nor its oxidized form, dehydroascorbate, can diffuse through membranes. Until now, two Asc transporters have been characterized in plants. AtDTX25, a member of the multidrug and toxic compound extrusion family in *A. thaliana*, is a vacuolar Asc transporter that controls intracellular iron cycling in seedlings (Hoang et al., 2021). AtPHT4;4 (AT4G00370.1), a member of the phosphate transporter 4 family of *A. thaliana* is found in the chloroplast envelope membrane (Guo et al., 2008; Miyaji et al., 2015; Fernie and Tóth, 2015). AtPHT4;4 knockout mutants (in the Ler-0 background) exhibited approximately a 30% decrease of chloroplastic Asc content, and accordingly, a slight impairment of the xanthophyll cycle and a decrease in NPQ at high light (Miyaji et al., 2015).

Ascorbate has been suggested to modify plant metabolism via distinct mechanisms. It has been suggested to maintain cellular redox homeostasis and participate in ROS signaling upon oxidative stress (Fotopoulos et al., 2006; Foyer et al., 2020). Ascorbate also serves as a chaperone for several 2-oxoglutarate-dependent dioxygenases (2-ODDs) participating in the biosynthesis of e.g., abscisic acid, gibberellins, and salicylic acid. Thereby, Asc was suggested to affect phytohormone-mediated signaling processes and modify stress tolerance and flowering time (Barth et al., 2006; Xiao et al., 2021). Based on the interconnections between the mitochondrial electron transport chain, Asc biosynthesis and photosynthesis, Asc was also proposed to take part in inter-organellar communication by acting as a metabolite signal (Rosado-Souza et al., 2020).

Chloroplastic Asc may play a central role in modulating metabolism since the chloroplast is a major sensor of environmental and metabolic changes, with its role in light and temperature acclimation well acknowledged (Kleine et al., 2021; Schwenkert et al., 2022). In addition, chloroplast-to-nucleus retrograde signaling adjusts gene expression and metabolism in order to maintain plant fitness (e.g., Leister, 2019).

In order to test the hypothesis that chloroplastic Asc may act as a metabolite signal, we compared the physiological and metabolic consequences of total cellular and chloroplastic Asc-deficiencies. To this end, we employed an Asc-biosynthesis mutant (*vtc2-4* in Col-0 background, Lim et al., 2016; Podmaniczki et al., 2021), and identified two novel PHT4;4 mutant lines in Col-0 background (*pht4;4-3* and *pht4;4-4*) to enable a faithful comparison of overall and chloroplastic deficiencies. Under moderate light intensities both Asc-deficiencies result in barely discernible effects on photosynthesis without signs of oxidative damage. On the other hand, we found widespread changes in the metabolome, involving remarkably similar metabolite sets upon total cellular and chloroplastic Asc-deficiencies. Finally, Proteome Integral Solubility Alteration (PISA) assay suggest that the modulating effect of Asc is exerted via interactions with specific chloroplastic proteins.

## Results

### Identification of new chloroplastic Asc-transporter mutants

The available and characterized Asc-deficient *VTC2* mutants are in Col-0 background (such as *vtc2-4*, used in this study), whereas the published *pht4;4-1* mutant is in Ler-0 background (Miyaji et al., 2015). In order to improve the comparability of overall and chloroplastic Asc-deficiencies, we identified two novel PHT4;4 mutants in Col-0 background that we named *pht4;4-3* and *pht4;4-4*. Homozygous T-DNA insertion mutants were identified via PCR and the exact positions of the T-DNA insertions were determined by sequencing (Fig. 1A). In the *pht4;4-3* mutant the insertion is found in the ninth intron, whereas in the *pht4;4-4* mutant it is located in the 5’UTR region.

**Figure 1.**
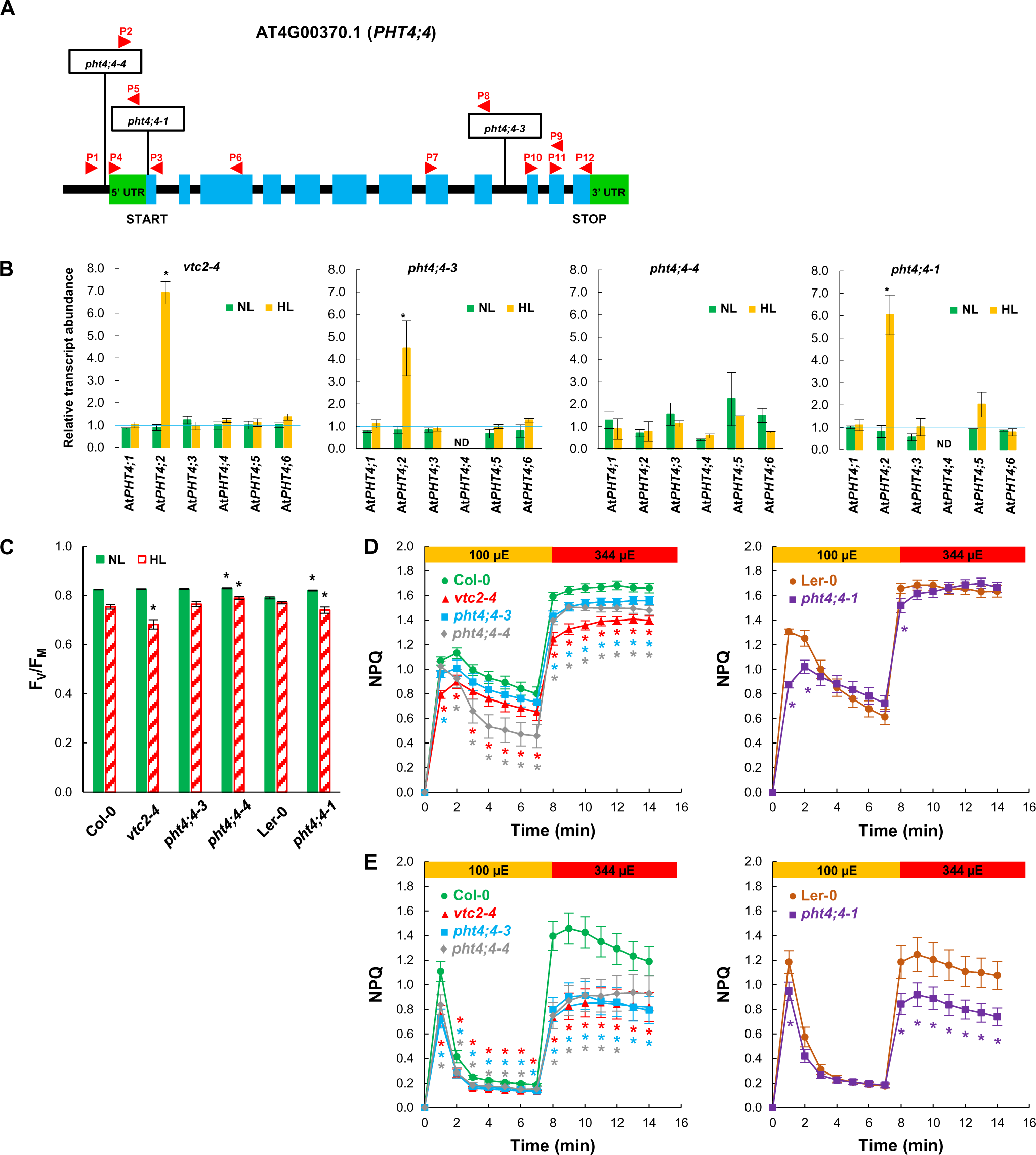
Characterization of *pht4;4* and *vtc2-4* mutants of *A. thaliana*. **A)** Physical map of *PHT4;4* (obtained from Phytozome, version 13) with the transposon and T-DNA insertion sites in the *pht4;4* mutants. Exons are shown in blue, and 5‘UTR and 3’UTR in green. The insertion sites are indicated by boxes, and the binding sites of the primers used for genotyping and gene expression analysis are shown as red arrows. **B)** Transcript levels of several *PHT4* genes as determined by RT-qPCR in *vtc2-4*, *pht4;4-3, pht4;4-4* and *pht4;4-*mutants relative to their respective WTs, at NL (approximately 80 μmol photons m^−2^s^−1^) or at HL (approximately 800 μmol photons m^−2^s^−1^ for two days), grown in three independent experiments. **C)** F_V_/F_M_ values of plants grown at NL (approximately 80 μmol photons m^−2^s^−1^) or at HL (approximately 800 μmol photons m^−2^s^−1^ for two days). The averages are based on three independent experiments, with nine replicates in each. **D)** Induction of NPQ at 100 and 344 μmol photons m^−2^s^−1^, of plants grown at NL. The averages are based on three independent experiments, with six biological replicates in each. **E)** Induction of NPQ at 100 and 344 μmol photons m^−2^s^−1^, of plants grown at HL. The averages are based on three independent experiments, with five replicates in each. Statistical significance levels between the mutants and their background strains were analyzed using Welch’s unpaired t-test. The significance levels are presented as *. Primers are listed in Suppl. Table 1.

The *PHT4;4* transcript, as determined by RT-PCR, was absent in the *pht4;4-1* and the *pht4;4-3* mutants, whereas it was moderately expressed in the *pht4;4-4* mutant (Suppl. Fig. 1A). *VTC2* transcript was detected at an equal level in all genotypes, except for the *vtc2-4* mutant, as expected (Suppl. Fig. 1A).

qRT-PCR analysis confirmed that the *PHT4;4* transcript was absent in the *pht4;4-1* and *pht4;4-3* mutants when grown at normal light (NL, 80 µmol photons m^−2^ s^−1^) and kept at high light intensity (HL, 800 µmol photons m^−2^ s^−1^) for two days. In the *pht4;4-4* mutant approximately 50% transcript abundance was observed relative to wild type (WT) at both light intensities (Fig. 1B, Suppl. File 1). The transcript levels of other PHT4 family members, which may possibly contribute to chloroplastic Asc transport, were also assessed. *PHT4;1, PHT4;3, PHT4;5,* and *PHT4;6* were mostly unaltered at both light conditions, whereas at HL, the transcript abundance of *PHT4;2* increased severalfold in the *pht4;4* knockout mutants and the *vtc2-4* mutant relative to their respective WT.

### The vtc2-4 and the pht4;4 mutants do not exhibit stress symptoms at normal light

Next, photosynthetic parameters were compared in plants kept under NL and HL conditions. The F_V_/F_M_ value, a widely used indicator for PSII photochemistry (Schansker et al., 2014; Stirbet et al., 2018), was similar (about 0.82) in all genotypes at NL, except for Ler-0, which displayed a slightly lower F_V_/F_M_ value (about 0.79, Fig. 1C). Following HL treatment, the F_V_/F_M_ value slightly decreased in all genotypes with the most substantial changes in the *vtc2-4* mutant.

In the following experiments, NPQ was assessed upon adaptation to 100 and 344 µmol photons m^−2^s^−1^ red light. As expected, NPQ was lower in the *vtc2-4* mutant than in Col-0 when grown at NL (in agreement with Müller-Moulé et al., 2002; Müller-Moulé et al., 2004) and NPQ was also diminished in the *pht4;4* mutants (Fig. 1D). Upon HL-treatment, the diminishment of NPQ was more enhanced in both the *vtc2-4* and the *pht4;4* mutants, especially when determined at 344 µmol photons m^−2^ s^−1^ red light (Fig. 1D,E).

The xanthophyll cycle pool (violaxanthin, antheraxanthin, and zeaxanthin) was about 20% smaller in the *vtc2-4* and *pht4;4* mutants relative to their WTs at NL. However, the pool size moderately increased upon the HL treatment in all genotypes (Fig. 2A, Suppl. File 1). The de-epoxidation indices were similar in the mutants and the wild types when grown at NL (Fig. 2B, Suppl. File 1). Upon HL treatment, the de-epoxidation index strongly increased; with augmentation being significantly less substantial in the Asc-deficient *vtc2-4* mutant and all *pht4;4* mutants compared to their respective WTs (Fig. 2B).

**Figure 2.**
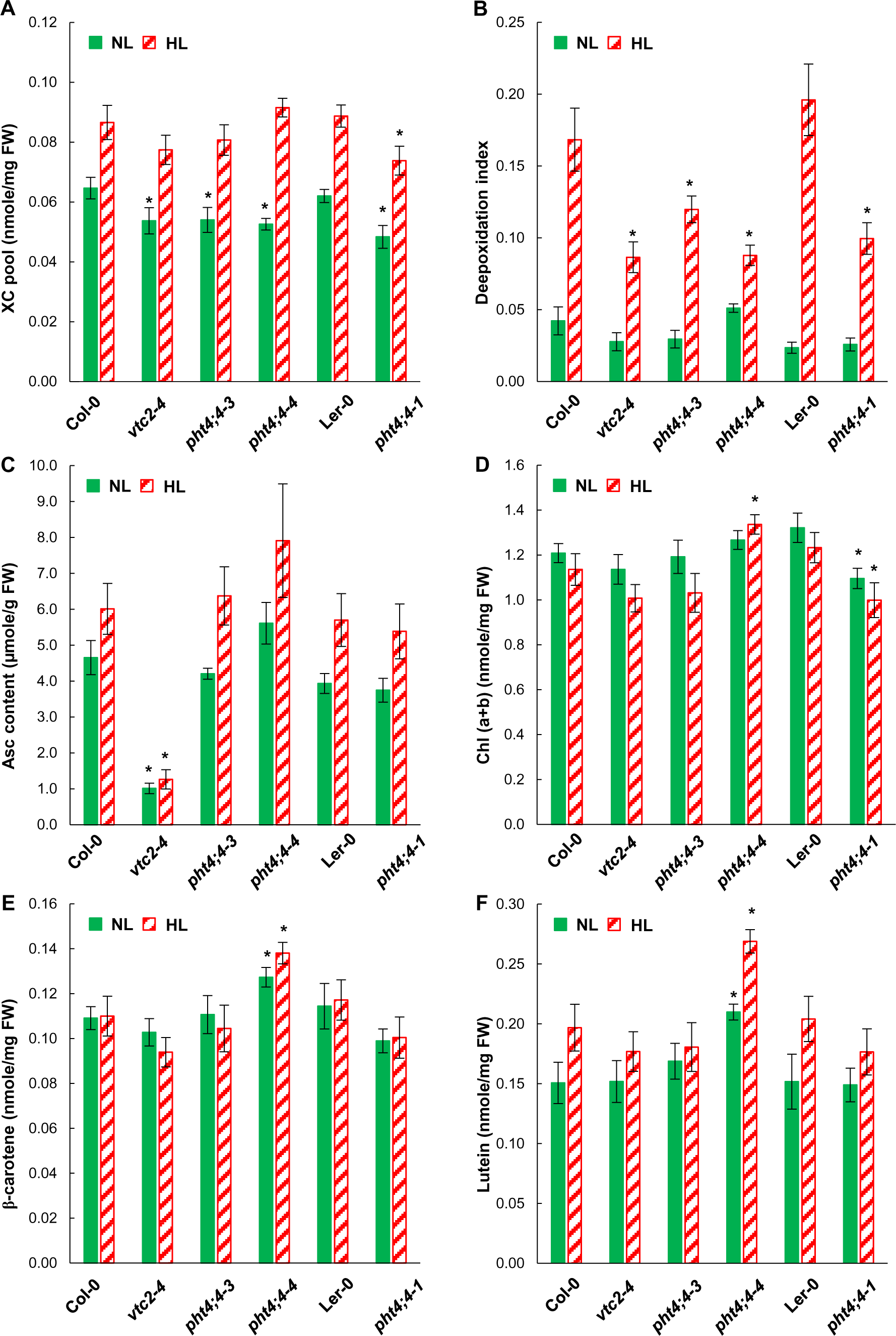
Carotenoids, Asc and Chl(a+b) contents in *pht4;4-1*, *pht4;4-3* and *vtc2-4* mutants and their respective WTs (Ler-0 and Col-0) grown at NL (80 μmol photons m^−2^s^−1^) or at HL (800 μmol photons m^−2^s^−1^ for two days). **A)** Xanthophyll cycle pigment pool size (violaxanthin, anteraxanthin and zeaxanthin, **B)** deepoxidation index, **C)** Asc content, **D)** Chl (a+b) content, **E)**, ß-carotene content, and **F)**, lutein content on a fresh weight basis. All the averages are based on three independent experiments with nine samples of each treatment and genotype. Statistical significance levels between the mutants and their background strains were analyzed using Welch’s unpaired t-test. The significance levels are presented as * p<0.1.

As expected, the total Asc content was low in the *vtc2-4* mutant (about 20% of Col-0, Fig. 2C; Lim et al., 2016). The Asc content of the *pht4;4* mutants were similar to their WT’s at NL. Upon HL treatment, an approximately 30 to 50% increase was observed in all genotypes, with the exception of the *vtc2-4* mutant (Fig. 2C, Suppl. File 1).

The Chl(a+b) contents were similar in all genotypes under NL conditions except for *pht4;4-1*, in which the Chl(a+b) content was significantly lower than in its WT (Fig. 2D, Suppl. File 1). There was a slight decrease in the Chl(a+b) content upon the HL treatment in all genotypes except the *pht4;4-4* mutant (Fig. 2D). The β-carotene contents were similar in all genotypes both under NL conditions and HL treatment with the exception of the *pht4;4-4* mutant that had a slightly higher β-carotene content (Fig. 2E, Suppl. File 1). The lutein contents of the various genotypes were also very similar under NL conditions, except for *pht4;4-4* in which it was slightly increased relative to Col-0. However, upon HL treatment the lutein content increased slightly (by 7 to 18% in *vtc2-4, pht4;4-3* and *pht4;4-1*) or moderately (by 28 to 34% in Col-0, *pht4;4-4* and Ler-0; Fig. 2F, Suppl. File 1).

In summary, the *vtc2-4* mutant and the newly identified *pht4;4* mutants have diminished NPQ and deepoxidation index but they do not exhibit any stress symptoms under NL conditions. Only the *vtc2-4* mutant showed a slightly enhanced stress sensitivity when exposed to HL treatment.

To assess the effects of Asc-deficiency on the plant metabolome, the *vtc2-4* and the *pht4;4-3* knockout mutants and their Col-0 background seemed to be the most suitable. In order to minimize environmental stress-related effects and to study the direct impact of total cellular and chloroplastic Asc-deficiencies, the plants were grown under NL conditions showing no phenotypic differences and stress symptoms, as described above.

### Assessment of chloroplastic Asc content

To assess the chloroplastic Asc contents of the *vtc2-4* and the *pht4;4-3* mutants and their Col-0 background, we first employed non-aqueous fractionation (NAF) to preserve chloroplastic Asc content (Krueger et al., 2011). Results indicate that the chloroplastic Asc level of the *pht4;4-3* mutant is approximately 60% lower than Col-0, while cytosolic Asc is about 100% higher, consistent with its role as an Asc-transporter. The *vtc2-4* mutant exhibited a drastic reduction, with approximately 95% lower chloroplastic Asc content than Col-0 (Suppl. Fig. 2A).

We corroborated results with Asc content determinations in isolated chloroplasts having at least 80% integrity (based on Aronsson and Jarvis, 2011; Joly and Carpentier, 2011). HPLC measurements revealed an approximately 35% decline in chloroplastic Asc level within the *pht4;4-3* mutant, and a decrease in the range of 90% for the *vtc2-4* mutant (Suppl. Fig. 2A).

Finally, we conducted *in vivo* Chl *a* fluorescence measurements (triggered by two 5-ms light pulses with varying time intervals) to evaluate the extent of chloroplastic Asc content reduction. This approach leverages the role of Asc as an alternative electron donor to PSII with heat-inactivated oxygen-evolving complexes, and has been shown to be sensitive to chloroplastic Asc content (Tóth et al., 2009). The regeneration kinetics of the F_20µs_/F_300µs_ parameter can be used as an estimate for the halftime (t_1/2_) of electron donation from Asc to PSII, specifically to Tyr ^+^ (Tóth et al., 2009).

Electron donation from Asc to PSII occurred with a t_1/2_ of approximately 20 ms in Col-0. It was slowest in the *vtc2-4* mutant (t_1/2_ approximately 36 ms), similar to the previously reported *vtc2-1* mutant (t_1/2_ approximately 34.6 ms, Suppl. Fig. 2C). The t_1/2_ in the *pht4;4-3* mutant was approximately 23.4 ms, confirming a milder reduction of chloroplastic Asc content compared to *vtc2-4*.

Thus, based on these three experimental approaches, we conclude an approximately 90% chloroplastic Asc content decrease in the *vtc2-4* mutant and a range of 30-50% for the *pht4;4* mutant compared to Col-0.

### Untargeted metabolomics reveals a global effect of chloroplastic Asc on the metabolome

Global metabolome analyses were performed on ion intensity data of metabolites annotated at two confidence levels. First, we annotated metabolites based on exact mass only (analogous with level “D” in Alseekh et al., 2021), yielding 234 putatively annotated metabolites that match compounds in the genome-scale metabolic network reconstruction of Arabidopsis (Dal’Molin et al., 2010). Within these, a subgroup of 118 metabolites could be further identified based on their mass spectrometric fragmentation patterns (analogous to level “B” in Alseekh et al., 2021; Suppl. File 2). The majority (>93%) of the metabolites showed a coefficient of variation of less than 40% across biological replicates, indicating high reproducibility.

Principal component analysis (PCA) revealed that the first two components almost entirely separated the three genotypes, and the analysis covered some 48% of the variance (Suppl. Fig. 3), indicating that the metabolome profiles are distinct. Size-proportional Venn diagrams represent genotype-specific and overlapping metabolite level differences that are significantly altered in the *vtc2-4* and *pht4;4-3* mutants relative to Col-0 (Fig. 3, Suppl. File 3). In the *vtc2-4* mutant, 81 and 66 metabolites were present at a significantly lower and higher level, respectively, relative to Col-0, which is, altogether, about 62% of all metabolites identified at level “D”. A remarkable proportion, approximately 40% of the metabolite changes overlapped between the Asc transporter (*pht4;4-3*) and biosynthesis (*vtc2-4*) mutants.

**Figure 3.**
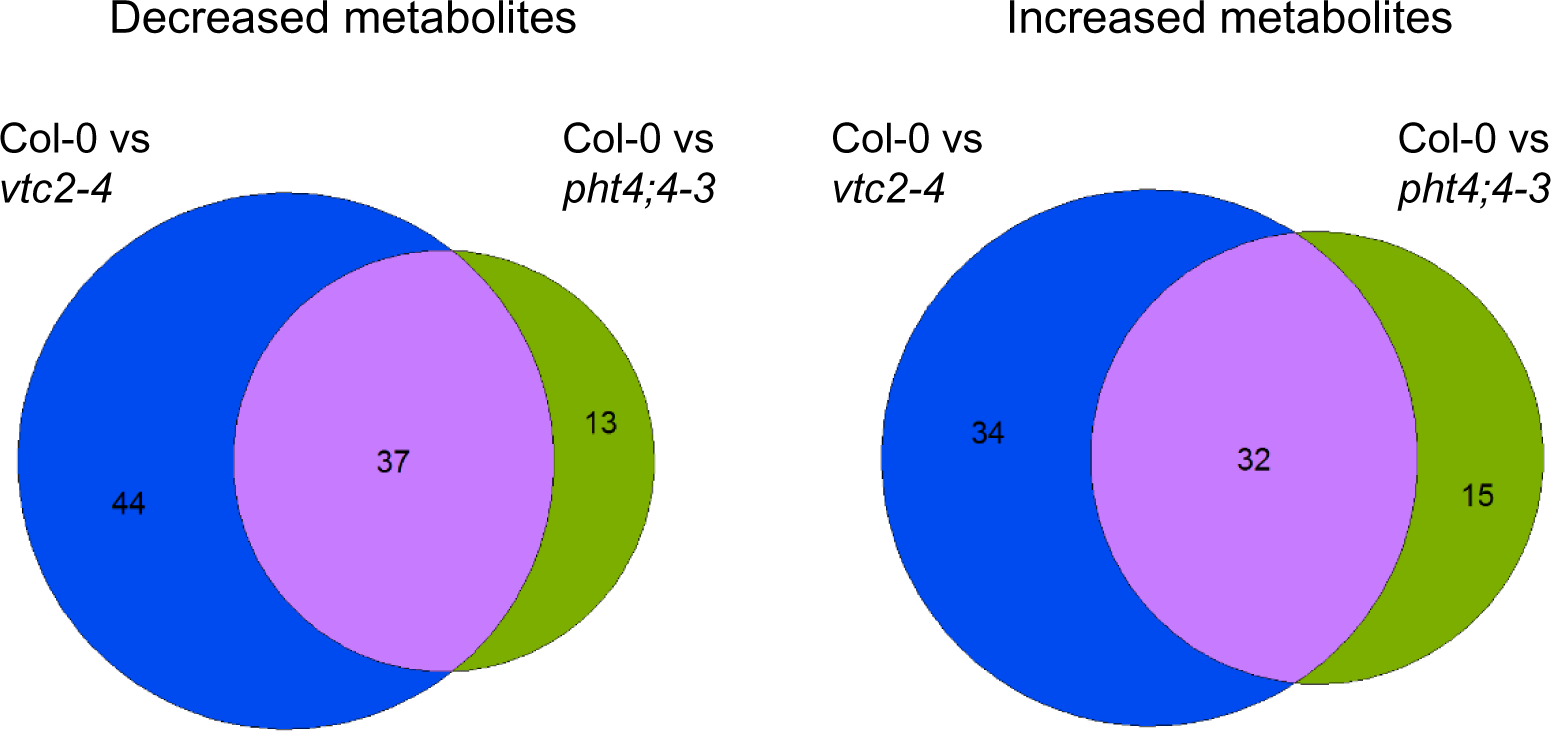
Size proportional Venn diagrams generated from metabolites annotated at levels “B” and “D” (see Materials and Methods). The diagrams represent genotype-specific and overlapping metabolite level differences that are significantly lower (**A**) or higher (**B**) in both genotypes relative to Col-0. Effect sizes are proportional to circle sizes. Blue colour represents *vtc2-4*-specific changes, green represents *pht4;4-3*-specific changes, and purple intersection range represent metabolite level changes that are significantly changed in both genotypes compared to Col-0.

We next analyzed the responses to Asc-deficiencies of the 118 individual metabolites annotated at level “B”. Of these metabolites, 67 and 42 were present at significantly different levels from those observed in Col-0 in *vtc2-4* and *pht4;4-3*, respectively (Fig. 4, Suppl. File 4). Of these, 34 metabolites were significantly altered compared to Col-0 in both the *pht4;4-3* and the *vtc2-4* mutants with the same change in direction (i.e., increase or decrease). This overlap is significantly higher compared than what we would expect just by chance based on the number of individual significant differences (randomization test p < 10^-5). Thus, these results show that Asc deficiency induces global metabolome alterations and this effect is largely mediated by chloroplastic Asc content.

**Figure 4.**
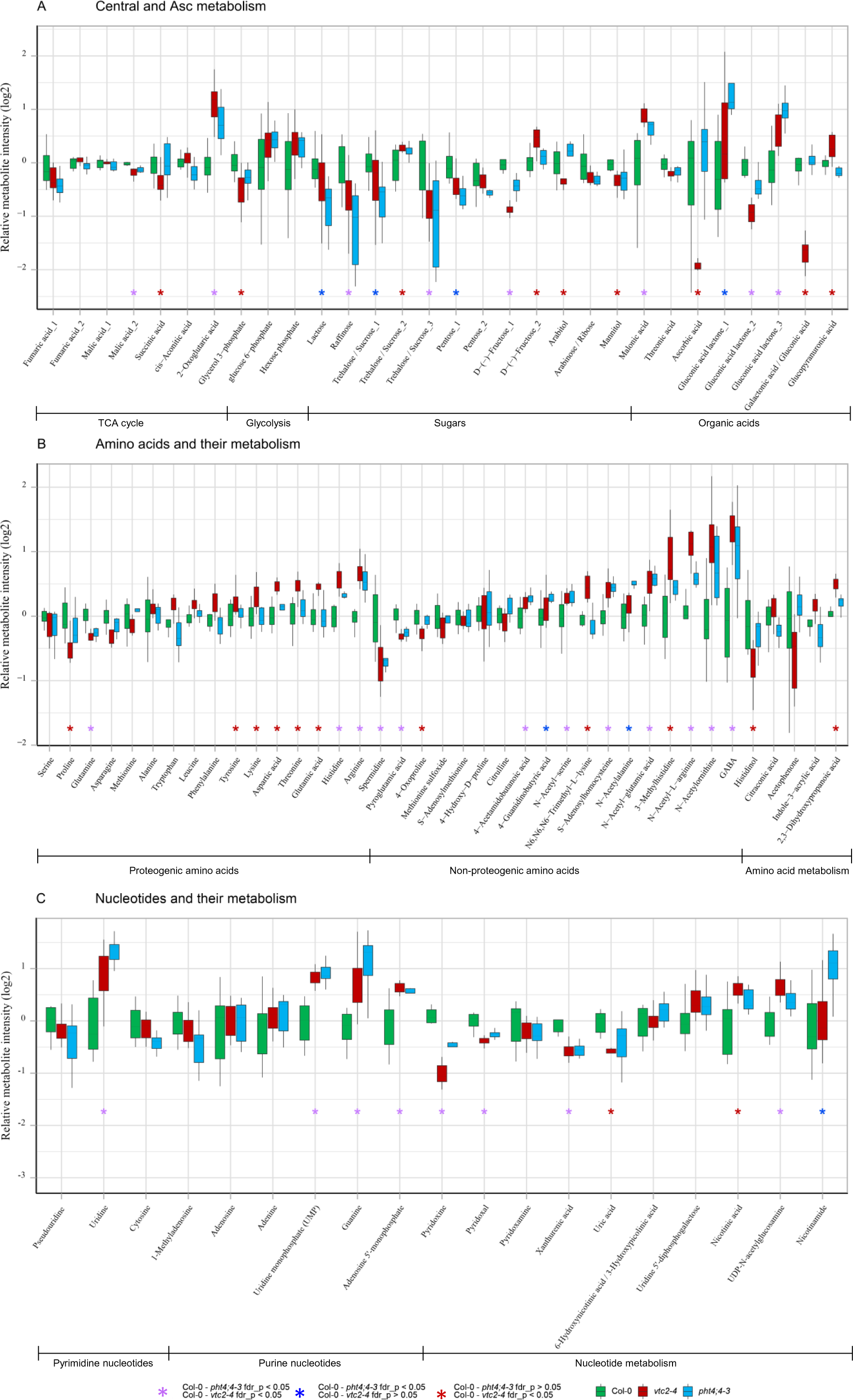
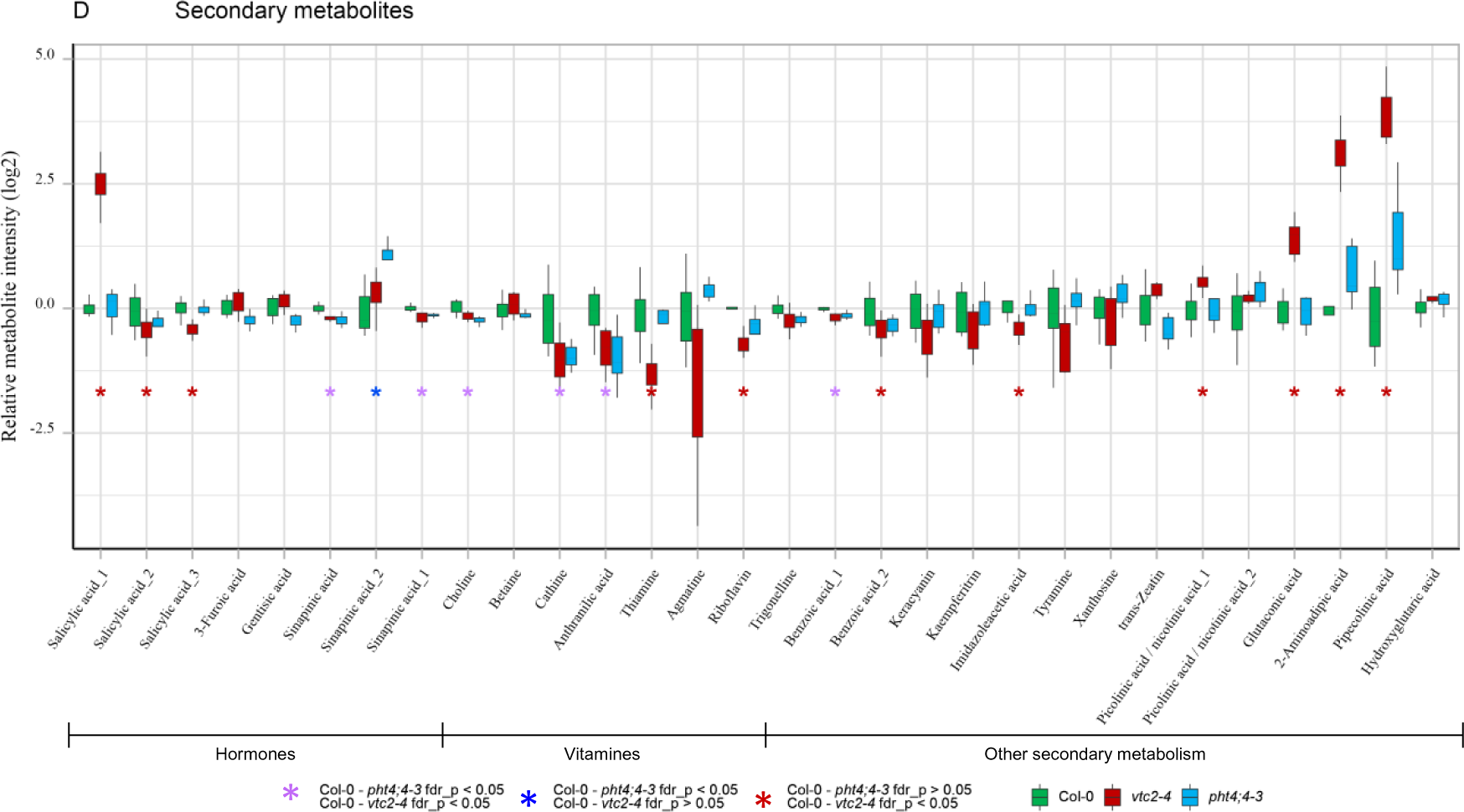
Metabolite levels annotated at level “B” (see Materials and Methods), in Col-0, *vtc2-4*, and *pht4;4-3* mutant lines. Box plots show median; interquartile range ±SEM fold change (log2 FC) to average levels in Col-0. Metabolites are grouped in four major compound classes, namely Central and Asc metabolism (**A**), Amino acids and their metabolism (**B**), Nucleotides and their metabolism (**C**) and secondary metabolism (**D**). All lines were measured in nine biological replicates in three cultivation batches. Statistics by t-test and multiple testing correction using “FDR” were used. Fold-changes, p-values, and adjusted p-values are listed in Suppl. File 4. Metabolites are grouped according to their functional classes. Colored “*” indicate the results of pairwise significance tests.

Next, Asc-deficiencies’ effects on the metabolites of the major compound classes were evaluated. A detailed description of the result can be found in Suppl. Text 1; here the major findings are summarized.

Regarding central and amino acid metabolism, we found a significantly increased level of the tricarboxylic acid (TCA) cycle intermediate, 2-oxoglutaric acid in both mutants (Fig. 4A), which may indicate increased respiration (Araújo et al., 2014, Dastogeer et al., 2017). In addition, arginine, a major nitrogen-storage compound in the plant cell (reviewed by Siddappa and Marathe, 2020), significantly increased in both mutants, and glutamate increased in *vtc2-4* (Fig. 4B). Along with this, the level of the plant signaling molecule γ-aminobutyric acid (GABA, Xu et al., 2021) and its derivatives, 4-acetamidobutanoic acid and 4-guaidinobutyric acid increased slightly in both mutants (Fig. 4B). This may indicate that the GABA shunt was activated to provide alternative carbon sources for mitochondrial respiration, similarly to e.g., salt stress (Che-Othman et al., 2020).

We observed that the amounts of proteinogenic amino acids (tyrosine, lysine, aspartic acid, threonine, and glutamic acid) increased mildly in the *vtc2-4* mutant (Fig. 4B), which may result from the activation of their biosynthetic pathways or enhanced protein degradation (Ishikawa et al., 2009, Lehmann et al., 2009, Hildebrandt, 2018). The levels of 2-aminoadipic acid and pipecolinic acid, which are both involved in lysine catabolism, have also significantly increased in the *vtc2-4* mutant (Fig. 4D). The levels of these amino acids remained essentially unchanged in the *pht4;4-3* mutant with the exception of histidine that increased in both mutants.

Chloroplastic Asc had a large effect on nucleotide metabolism, too (Fig. 4C). The increase in uridine and uridine monophosphate levels, and in malonic acid levels (Fig. 4A) indicates activation of the pyrimidine (uridine) salvage pathway that plays crucial roles in photoassimilate allocation and partitioning (Chen and Thelen 2011). Nicotinic acid levels also increased in both mutants, and the level of nicotinamide was significantly higher in the *pht4;4-3* mutant than in Col-0 (Fig. 4C). These compounds may be re-utilised for the synthesis of pyridine nucleotides by salvage pathways and the synthesis of pyridine alkaloids (Ashihara et al., 2015, Gakière et al., 2018).

We also obtained data on stress-responsive metabolites. The amounts of osmolyte sugars that are known to accumulate under oxidative stress conditions, such as raffinose, fructose, trehalose/sucrose, and mannitol (Baxter et al., 2007, Lehmann et al., 2009) remained unaltered or changed only mildly (Fig. 4A). The amounts of amino acids with antioxidant and osmolyte properties (proline, citrulline, spermidine) also remained unchanged or decreased (Fig. 4B). The amounts of uridine, uridine monophosphate, guanine, and adenosine 5’-monophosphate significantly increased in both mutants (Fig. 4C). Previous studies established that nucleotide metabolism and synthesis are frequently diminished upon oxidative stress (Baxter et al., 2007). Uric acid has been observed to decrease under oxidative stress conditions (Sipari et al., 2020); here we observed a significant decrease in uric acid level only in the *vtc2-4* mutant (Fig. 4C). The thiamine level slightly decreased in the *vtc2-4* and it remained at a similar level in the *pht4;4-3* mutant (Fig. 4D). Thiamine functions as an important stress-response molecule that alleviates oxidative stress during different abiotic stress conditions (Tunc-Ozdemir et al., 2009; Rosado-Souza et al., 2020). Similarly, pyridoxine (vitamin B6), and its vitamer derivatives, pyridoxal, and pyridoxamine, have been shown to act as antioxidants and their levels increase strongly upon photooxidative stress (Chen and Xiong 2005; Havaux et al., 2009; Parra et al., 2018). Here, we observed a significant decrease in pyridoxine and pyridoxal levels in both mutants (Fig. 4C).

Thus, these results show a global and extensive response of cellular metabolism to the diminishment of total cellular and chloroplastic Asc contents. The most remarkable changes were the activation of the GABA shunt and the upregulation of nucleotide salvage pathways. The considerable overlap between the two *vtc2-4* and the *pht4;4-3* mutants suggests these effects were exerted through chloroplastic Asc content. On the other hand, changes specific to the *vtc2-4* mutant were observed regarding amino acid metabolism. Importantly, no signs of oxidative stress were observed in either mutant, in agreement with the photosynthesis results (Figs. 1 and 2).

### Proteome Integral Solubility Alteration (PISA) assay reveals interaction between Asc and chloroplastic proteins

Recent advancements in tandem mass spectrometry (MS) have enabled large-scale analysis of protein structure and conformation. These techniques can determine the thermal denaturation profile of hundreds to thousands of proteins simultaneously, providing insights into protein-protein interactions and other factors influencing protein stability (Volkening et al., 2019).

One approach to elucidate the influence of interacting molecules on protein function involves manipulating extrinsic factors, such as temperature, to induce controlled alterations in protein properties. In thermal proteome profiling (TPP), protein stability is monitored through changes in solubility at different temperatures (Savitski et al., 2014; Mateus et al., 2020). Interaction with certain metabolites can alter protein stability, either directly through binding or indirectly through changes in protein folding. For example, Sridharan et al. (2019) demonstrated this with ATP and GTP.

Recently, a high-throughput version of TPP was developed, called Proteome Integral Solubility Alteration (PISA) assay, which increases the analysis throughput by one to two orders of magnitude (Gaetani et al., 2019). In the PISA assay, a sample is subjected to a treatment expected to induce conformational changes, then aliquoted and treated at different temperatures. The increased throughput of the PISA assay compared to regular TPP is achieved through pooling of protein aliqots after individual temperature treatments, reducing the number of analyzed samples. The abundance of the pooled soluble fraction reflects the integral of the melting curve, and comparison with a reference sample allows measurement of relative changes in thermal stability. Unlike conventional TPP, the PISA assay avoids fitting melting curves to the data, mitigating potential bias for proteins with non-standard melting profiles (Gaetani et al., 2019).

Using the PISA assay, we aimed to test whether Asc interacts with chloroplastic proteins. To this end, we profiled the thermal stability of over 600 chloroplastic Arabidopsis proteins, in the presence of 0, 2, 5 and 10 mM Asc (Suppl. Files 5 and 6). The employed Asc concentrations are physiologically realistic, as the chloroplastic Asc concentration is in the range of 10 mM in vascular plants (Foyer and Lelandais, 1995).

To investigate the stability of Asc during sample preparation for the PISA assay, which involves a 3-minute treatment at 40-55 °C, we assessed the levels of Asc, DHA, and potential degradation products (2-keto-gulonic acid and L-tartaric acid) using HPLC and MS analyses. Suppl. Fig. 4A shows that the Asc content in the chloroplast isolate matched the added 10 mM Asc and remained unchanged following heat treatment at 55 °C. DHA levels were minimal, ranging from 3 to 4% with or without heat treatment. 2-keto-gulonic acid and L-tartaric acid were found in the μM concentration range, corresponding to a total amount of 0.26% and 0.22% in the room temperature and high temperature-incubated PISA samples, respectively (Suppl. Fig. 4B). These findings suggest that the degradation compounds of Asc are unlikely to affect the PISA assay.

Altered thermostability for numerous chloroplastic proteins was observed in the presence of 10 mM Asc. At an adj. p-value treshold of 0.05, 69 interacting proteins were found, which are involved in central carbon metabolism (17.6%), amino acid metabolism (22.1%), redox regulation and stress response (17.6%), photosynthetic electron transport (7.4%) and other processes or with unknown function (35.3%) (Fig. 5A, Suppl. File 5). Thus, the list of putatively interacting proteins covers a wide range of biological processes, suggesting that Asc has a high potential to modify enzyme activities and/or protein structures.

**Figure 5.**
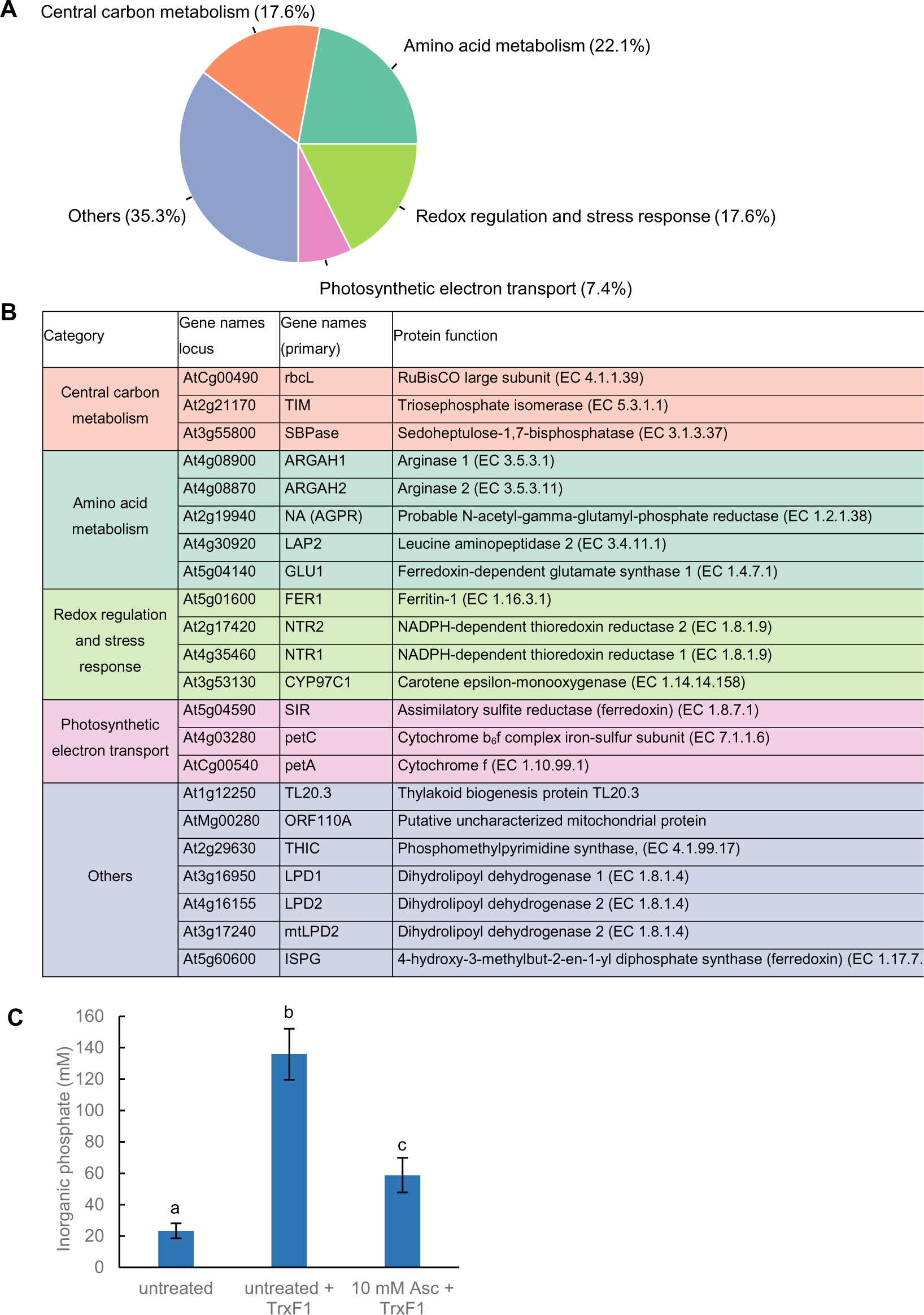
Chloroplastic proteins with altered thermostability (adj. p-value < 0.05) in response to 5 and 10 mM Asc based on the PISA assay. Proteins were grouped based on the chosen biological processes: photosynthetic electron transport, central carbon metabolism, redox regulation and stress response, and amino acid metabolism. Central carbon metabolism included glycolysis, the pentose phosphate pathway, the tricarboxylic acid cycle cycle, carbon fixation, fatty acid metabolism and related sugar metabolism. **A**) Classification of major protein groups in response to 10 mM represented as a pie chart **B**) The list of proteins with altered thermostability in response to 5 mM Asc (see Suppl. File 5). **C)** *In vitro* activity of SBPase in the presence of 10 mM Asc. SBPase and thioredoxin F1 were expressed in recombinantly in *E. coli* and purified. The assay measures the conversion rate of the alternative SBPase substrate fructose-1-6-bisphosphate to fructose-6-phosphate through the accumulation of inorganic phosphate. The significance of differences between means were determined by ANOVA with Tukey post-hoc test. The means with different letters are significantly different (P < 0.01).

In the search for high-affinity interactions, we assessed the effects of a lower Asc concentration, 5 mM, on the thermostability of chloroplastic proteins (Fig. 5B). At an adj. p-value < 0.05, 22 hits were found, of which the majority, 14 and 17 were found in the presence of 10 mM Asc at an adj. p-value < 0.05 and p-value < 0.1, respectively (Suppl. File 5).

The thermostability of several Calvin-Benson cycle enzymes was affected by Asc. These include RuBisCO large subunit (rbcL), sedoheptulose-1,7-bisphosphatase (SBPase), and triosephosphate isomerase (TIM; Fig. 5B). It has been proposed that the Calvin-Benson cycle enzymes interact within a multienzyme complex (Suss et al., 1993, Graciet et al., 2004, Yu et al., 2020), in which the enzymes are activated under reducing conditions, for instance, by the thioredoxin system (Marri et al., 2009; Kang et al., 2019). We hypothesize that Asc may modify these reactions.

Other possible interacting partners of Asc are arginase 1 (ARGAH1) and arginase 2 (ARGAH2). These enzymes are predicted to be localized to the chloroplasts and mitochondria and responsible for the degradation of arginine (Patel et al., 2017; Siddappa and Marathe, 2020). The reaction catalyzed by arginase yields ornithine which is the precursor of polyamines, proline and glutamate that participate in defense against oxidative stress. In addition, the thermostability of At2g19940, which is a putative chloroplast localized N-acetyl-glutamatyl-phosphate reductase (AGPR) participating in ornithine biosynthesis (Slocum, 2005; Winter et al., 2015), was also affected by Asc (Fig. 5B). In line with these results, the metabolomics analysis (Fig. 4) show that arginine accumulates, whereas proline, glutathione and spermidine levels decrease suggesting that arginase activity was downregulated upon Asc-deficiency. In relation to this, the thermostability of leucine aminopeptidase 2 (LAP2) was significantly altered by Asc. LAP2 liberates N-terminal leucine, methionine and phenyl alanine from proteins and peptides thereby it controls amino acid turnover. Nitrogen-rich amino acids, such as glutamate and glutamine, and GABA levels are particularly affected by LAP2 activity (Waditee-Sirisattha et al., 2011). The effect of Asc on LAP2 thermostability is in line with the changes in GABA content upon Asc-deficiency (Fig. 4).

Asc affected the thermostability of ferredoxin-dependent glutamate synthase 1 (GLU) as well, found both in the chloroplast and the mitochondria (Jamai et al., 2009). GLU is responsible for the reassimilation of photorespiratory ammonia as well as for primary nitrogen assimilation in the chloroplast. In this reaction, glutamate is produced from glutamine and 2-oxoglutarate, generated in the TCA cycle (Yoneyama and Suzuki, 2020). In broad agreement with the PISA assay, metabolomics showed enhanced glutamate content in the *vtc2-4* Asc-deficient mutant (Fig. 4).

Another potential interacting partner is chloroplastic assimilatory sulfite reductase (SIR, Fig. 5B) that is essential for plant survival due to its role in assimilatory sulfate reduction. Its downregulation causes severe adaptive reactions of primary and secondary metabolism (Khan et al., 2010, Wang et al., 2019). SIR receives electrons from ferredoxin at the acceptor side of PSI and it reduces sulfite (SO_3_^2-^) to hydrogen sulfide (H_2_S). This step is followed by the incorporation of S_2_^−^ into the amino acid skeleton of the intermediate O-acetyl-serine (Feldman-Salit et al., 2019) and the generation of cysteine. Besides this reaction, H_2_S also acts as a signaling molecule thereby modifies plant metabolism at multiple levels (Thakur and Anand, 2021). The regulation of SIR is still elusive, but must be an integral part of the coordination of the distribution of reducing power from PSI via ferredoxins to the various other consumers (Telman and Dietz, 2019).

Asc also affected proteins involved in the regulation of photosynthetic electron transport. We found that thermostability of two subunits of the cytochrome (Cyt) b_6_f complex, namely PetA (Cytf) and PetC (the Rieske iron-sulfur protein) were significantly influenced by the Asc treatment (Fig. 5B). The Cytb_6_f complex is known to be particularly suited to sensing the redox state of the electron transport chain and the chloroplast stroma (reviewed by Malone et al., 2021). In agreement with the PISA assay, it was shown earlier that Asc reduces both PetA and PetC in the low mM range (Riedel et al., 1991, Zhang et al., 2001, Malone et al., 2019). A detailed description of the remaining putative interactions can be found in Suppl. Text 2.

Therefore, several lines of evidence suggest that Asc-protein interactions cause at least some of the observed metabolomic changes. To investigate this further, we focused on SBPase, a one of the top candidate affected by Asc according to the PISA assay. We recombinantly expressed and purified SBPase in E. coli, along with its known activator, thioredoxin F1 (Gütle et al., 2016). Remarkably, we found a significant decrease (approximately 57%) in the *in vitro* activity of SBPase upon incubation with 10 mM Asc (Suppl. Fig. 5). This finding highlight a direct inhibitory effect of Asc on SBPase activity, potentially contributing to the observed metabolomic changes.

## Discussion

It has been hypothesized earlier that chloroplastic Asc may modulate plant metabolism by acting as a metabolite signal (Rosado-Souza et al., 2020). We took two complementary approaches to test this hypothesis. First, we comprehensively compared the physiological and metabolomic effects of total cellular and chloroplastic Asc deficiencies by employing an Asc-biosynthesis mutant (*vtc2-4;* Lim et al., 2016), and a novel knock-out mutant of the PHT4;4 transporter in Col-0 background (*pht4;4-3*). Second, we used proteome-wide thermostability analysis to reveal potential interactions between Asc and chloroplastic proteins.

The extent of chloroplastic Asc decrease is in the range of 90% in *vtc2* mutants (Suppl. Fig. 2). Asc is required for violaxanthin de-epoxidase activity (Bratt et al., 1995; Saga et al., 2010; Hallin et al., 2016); accordingly, Asc-deficiency diminishes NPQ, particularly in HL (Fig. 1) and the de-epoxidation index was approximately 50% lower in the *vtc2-4* mutant than in Col-0 in HL (Fig. 2).

In the *pht4;4* mutants, the total cellular Asc concentration is unaltered relative to the WTs (Fig. 2C). We found that the chloroplasts of the *pht4;4-3* mutant contain about 30-50% less Asc than WT chloroplasts (Suppl. Fig. 2). In agreement with this, there was a discernible decrease in NPQ and in the de-epoxidation index in the *pht4;4* mutants relative to their WT (Figs. 1, 2).

Ascorbate is recognized as one of the most important antioxidants in the cell. In this respect, it is surprising that Asc-deficient mutants have the same levels of β-carotene and lutein as the WT under NL conditions (Fig. 2). The amounts of osmolyte sugars (e.g., raffinose, fructose, mannitol) and amino acids (proline, citrulline, spermidine) participating in oxidative stress response also remained or changed only mildly (Fig. 4). In agreement with our data, it has been observed earlier that under regular laboratory conditions, glutathione, Asc peroxidase activity, and the redox state of Asc are also unaltered in the *vtc2* mutants, just as well as photosynthetic activity (Müller-Moulé et al., 2004; Lim et al., 2016, Fig. 2), and only moderate changes occur at constant HL (Müller-Moulé et al, 2004, Figs. 1, 2).

Thus, an approx. 90% reduction of total cellular Asc content in the *vtc2-4* mutant or the 30-50% chloroplastic Asc content decrease in the *pht4;4-3* mutant did not lead to photooxidative stress under NL conditions, indicating a margin of safety in Asc levels under such conditions. We note that the partial decrease in chloroplastic Asc content of the *pht4;4-3* mutant suggest that Arabidopsis may possess other, hitherto unknown chloroplastic Asc transporters. A possible candidate may be PHT4;2, whose expression level increased severalfold upon HL treatment (Fig. 1); future studies should test this possibility.

To reveal how cellular metabolism is altered upon total cellular and chloroplastic Asc deficiencies, untargeted metabolomics was employed on Col-0, the *vtc2-4* and the *pht4;4-3* knockout mutants that are both in Col-0 background. In order to minimize environmental stress-related effects on the metabolome, the plants were grown under precisely controlled NL conditions.

Both total cellular and chloroplastic deficiencies caused large and widespread changes in the metabolome, with the metabolites synthesized both in the chloroplast and the cytosol. The changes affected approximately 57% of the annotated compounds in the *vtc2-4* mutant of which about 51% occurred in the *pht4;4-3* mutant as well (Fig. 4). The fact that the metabolite changes largely overlapped between the mutants, suggests that chloroplastic Asc content has a particularly important role in modulating metabolic pathways. In the search for a possible mechanism on how Asc could affect the metabolome, the PISA assay revealed that Asc has a potential to interact with various chloroplastic proteins, participating in arginine metabolism, photosynthetic electron transport and CO_2_ fixation.

Members of the TCA cycle were mildly affected by Asc-deficiency, with the exception of 2-oxoglutarate being significantly increased in both mutants. Along with this, the levels of GABA, its derivatives and arginine significantly increased in both mutants, and glutamate increased in the *vtc2-4* mutant (see Fig. 6 for a summarizing figure), suggesting that the GABA shunt is activated upon chloroplastic Asc-deficiency. In line with the increase in arginine content, we found putative interactions between Asc and ARGAH1 / ARGAH2, participating in arginine degradation (Patel et al., 2017; Siddappa and Marathe, 2020). In relation to this, LAP2, controlling amino acid turnover, glutamate, glutamine, and GABA levels (Waditee-Sirisattha et al., 2011), was affected by Asc. In addition, the PISA assay also revealed a possible interaction between Asc and GLU, producing glutamate from glutamine and 2-oxoglutarate in the chloroplast (Yoneyama and Suzuki, 2020). Thus Asc may interact with several enzymes regulating the GABA shunt. The main role of the GABA shunt is to provide alternative carbon sources for mitochondrial respiration; GABA has also been described to participate in signal transduction and modulate metabolism in various ways (Xu et al., 2021), therefore, it may potentially transmit the effects of Asc-deficiency to various other metabolic pathways. We noted that despite the increase in Glu levels, proline and the polyamine spermidine decreased in both mutants (see Fig. 6).

**Figure 6.**
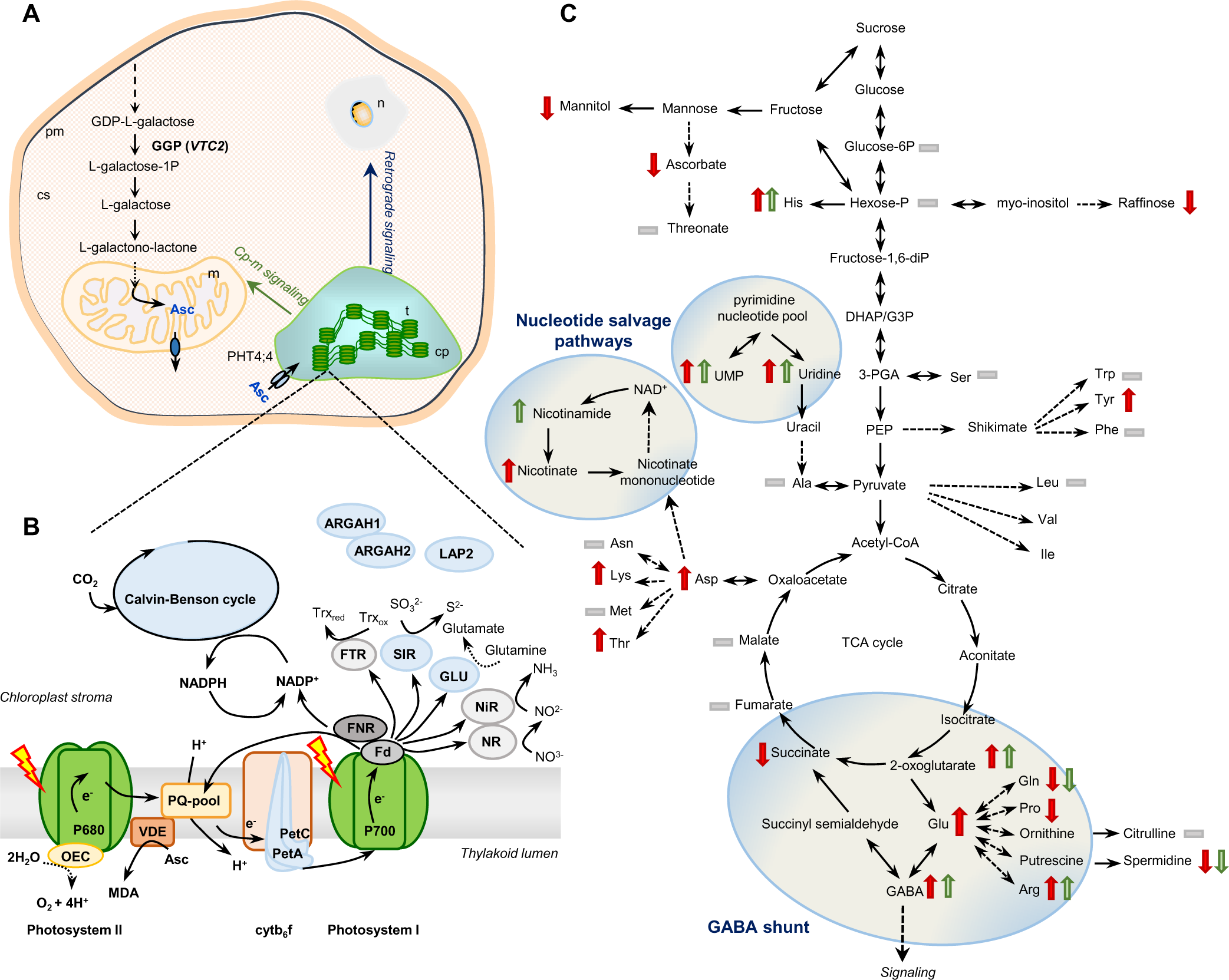
Chloroplastic Asc is a regulatory hub in plant metabolism **A)** Asc biosynthesis takes place in the cytosol, with the exception of the last step, occurring in mitochondria. *vtc2* encodes GDP-L-galactose phosphorylase (GGP), catalyzing the first committed step of Asc biosynthesis. PHT4;4 transports Asc into the chloroplast. **B)** Photosynthetic electron transport components and stromal enzymes potentially interacting with chloroplastic Asc (indicated in blue, based on the PISA assay of this study). **C)** Chloroplastic and cellular Asc-deficiencies lead to widespread changes in the metabolome, including the induction of the GABA shunt and nucleotide salvage pathways (circled). Red and green arrows indicate significant changes in the respective metabolites identified at level “B” (see Fig. 4) in the *vtc2-4* and the *pht4;4-3* mutants compared to the WT (Col-0), respectively. Grey bars mean that the given metabolite was identified but no significant changes relative to the WT were detected.

Cellular and chloroplastic Asc-deficiencies also resulted in the induction of nucleotide salvage pathways, probably to fine-tune photoassimilate allocation and partitioning. We note that induction of the GABA shunt along with nicotinamide and uridine salvage pathways was observed earlier upon aging (Schippers et al., 2008, Sipari et al., 2020).

We found that Asc may also affect SIR activity; SIR is another important regulatory enzyme participating in adaptive reactions of primary and secondary metabolism and signaling (Thakur and Anand, 2021).

The PISA assay revealed that Asc has the potential to interact with Calvin-Benson cycle enzymes as well, including SBPase. Using *in vitro* assays, we directly demonstrated Asc’s inhibitory effect on SBPase activity (Suppl. Fig. 5). This may be particularly interesting, considering that SBPase activity is associated with growth, starch accumulation, and chilling stress tolerance (Ding et al., 2017; Liu et al., 2012). We note that metabolite-level regulation of the Calvin-Benson cycle activity has been also described in cyanobacteria (Sporre et al., 2022).

Finally, Asc was found to interact with two cytb_6_f components, PetA and PetB. A direct effect of this interaction on the metabolome could not be identified in this study. Nevertheless, we note that the activity of cytb_6_f controls the redox state of the photosynthetic electron transport (Schöttler and Tóth, 2015), which is a major retrograde signal, affecting the expression of a large number of genes (Gjindali and Johnson, 2023). Therefore, by interacting with PetA and PetB, Asc may participate in retrograde signaling, thereby modulating plant metabolism.

Thus, our results support the earlier hypothesis by Rosado-Souza et al. (2020) and Kleine et al. (2021) that Asc may eventually act as a metabolite signal to mediate metabolic crosstalk between the chloroplast and mitochondria, and possibly as a retrograde signal to modify the expression of nuclear genes. As sizeable oxidative stress did not occur, the observed modulating effect of Asc is likely to be unrelated to ROS-signaling pathways, although, we do not exclude the possibility that under severe stress conditions, the ROS-signaling pathways would also be affected by Asc-deficiency (Foyer et al., 2020).

A subset of metabolites changed significantly in the *vtc2-4* mutant only (Suppl. Table 2), indicating that these changes are linked to cytosolic Asc content. These include free amino acids, such as tyrosine, lysine, aspartic acid, threonine, and glutamic acid and secondary metabolites, among which salicylic acid and thiamine have been shown earlier to interact with Asc metabolism and function (Nazar et al., 2015; Rosado-Souza et al., 2020). The synthesis of salicylic acid and thiamine takes place in the cytosol. Thus it can be assumed that their altered biosynthesis is a consequence of the diminished cytosolic Asc content and/or Asc biosynthesis intermediates. On the other hand, it is also to be considered that in the *pht4;4-3* mutant the reduction in chloroplastic Asc concentration is somewhat lower than in the *vtc2-4* mutant thus the effect on the alteration on metabolites may be milder.

As discussed above, substantial oxidative stress was unlikely to occur in the mutants under NL conditions. Therefore, the differences in metabolite levels are not due to oxidative stress but may reflect a specific response to decreased chloroplastic Asc contents. In nature, chloroplastic Asc-deficiency may occur under low light conditions, since Asc biosynthesis is known to be strongly dependent on light and the photosynthetic electron transport in particular (Yabuta et al., 2007; Bartoli et al., 2009). Low photosynthetic rate occurs under various stress conditions, namely in low temperature (Crosatti et al., 2013), excessive illumination (Gururani et al., 2015), drought stress (Feller, 2016), and heat stress (Yamamoto, 2016) that may all limit the rate of Asc biosynthesis and, at the same time, enhance Asc oxidation. All these conditions require adaptive responses and our data suggest that a major player in environmental stress adaptation is Asc, acting as a metabolite signal.

Overall, our study uncovered that lowering chloroplastic Asc levels triggers a global rewiring of plant metabolism, suggesting a novel regulatory mechanism. This rewiring may may involve interactions between chloroplastic Asc and proteins governing photosynthesis, cellular regulation, and potentially, retrograde signaling. These findings raise the intriguing possibility that plants utilize chloroplastic Asc as a sensor to perceive environmental challenges and adapt their metabolism accordingly.

## Materials and methods

### Plant material and growth conditions

The Asc-deficient *vtc2-4* mutant in the Col-0 background (TAIR stock No: CS69540; see Lim et al., 2016) originates from the laboratory of Dr. John F. Golz (University of Melbourne, Australia). The *pht4;4-1* mutant (with Ler-0 backgound, Miyaji et al., 2015) was purchased from TAIR (stock No: CS26443). Two novel AtPHT4;4 mutants in Col-0 background were identified and characterized (TAIR stock No: CS469134 and CS444342), which we named *pht4;4-3* and *pht4;4-4*.

All *Arabidopsis thaliana* genotypes were initially grown in a growth chamber under short-day conditions (8 h light, 16 h dark), at approximately 80 µmol photons m^−2^ s^−1^ in the light period (low light, NL) for eight weeks. The temperature was 18°C in the dark and 22°C in the light. When the plants were eight weeks old, a subset was transferred to high light (HL) (approximately 800 µmol photons m^−2^s^−1^) for two days.

We noted that the Asc-deficient T-DNA insertional mutant *vtc2-4* does not show any alteration in the phenotype in comparison with Col-0 when grown under short-day conditions (in agreement with Lim et al., 2016). The phenotype of the newly characterized *pht4;4-3* and *pht4;4-4* mutants were also indistinguishable from their Col-0 background. On the other hand, the *pht4;4-1* mutant plants had long petioles relative to Ler-0, a difference that was not observed by Miyaji et al., (2015) when studying four-week-old plants grown under long-day conditions (Suppl. Fig. 1B,C).

### PCR, RT-PCR and RT-qPCR reactions

DNA was isolated based on the protocol of Edwards et al. (1991) and the T-DNA insertion in the *pht4;4-3* and *pht4;4-4* mutants was confirmed by PCR, using gene-specific forward, reverse and T-DNA-specific primers, respectively (see Suppl. Table 1 for the list of primers).

For RNA isolation, approximately 50-100 mg of leaf material were collected and frozen in liquid nitrogen before grinding with a MM400 laboratory mill (Retsch, Germany) at 30 Hz for 1 min. Direct-Zol RNA isolation kit was used, following the recommendation of the manufacturer (Zymo Research). RNA integrity was checked on a 1% (w/v) denaturing agarose gel. To remove contaminating DNA from the samples, the isolated RNA was treated with DNaseI (Zymo Research). 1 µg of total RNA was used for cDNA synthesis with random hexamers using FIRESript reverse transcriptase, following the recommendation of the manufacturer (Solis BioDyne). To confirm the absence of transcript in the mutant lines, RT-PCR assays were conducted, consisting of 30 amplification cycles using primers listed in Suppl. Table 1.

The relative transcript abundance of several *PHT4* genes was determined via qRT-PCR following the method described in Vidal-Meireles et al. (2017). The RT-qPCR data are presented as the fold change in mRNA transcript abundance normalized to the average of three reference genes (*GAPDH, ACT2, UBC21*) and relative to the respective WT strains (see Suppl. Table 1 for the list of primers).

### Ascorbate, chlorophyll and carotenoid content determination

Total Asc contents of leaves (20-40 mg of each sample), and Asc and DHA contents of intact chloroplasts were determined by HPLC, based on the protocol of Kovács et al. (2016).

The assessment of chloroplastic Asc content was performed by non-aqueous fractionation (NAF). For this, Arabidopsis leaves were harvested and snap-frozen in liquid nitrogen. Two grams of fresh weight of frozen plant material was ground to a fine powder and freeze-dried at 0.02 bar for 5 days in a lyophilizer, which had been pre-cooled to −40°C. The NAF-fractionation procedure was performed as described in Shapiguzov et al., (2019) and Medeiros et al (2019). Subcellular compartmentation of markers and the metabolites of our interest was calculated by the BestFit method as described in Krueger et al., (2011), Klie et al., (2011), Krueger et al., (2014). BestFit applied the linear regressions for subcellular compartments using the percentage of distribution of each metabolite of our interest and the subcellular markers of each fraction. Marker measurements for NAF have been performed as described in Shapiguzov et al., (2019) and Medeiros et al (2019).

For carotenoid and Chl(a+b) determination, approximately 10-20 mg of Arabidopsis leaves were frozen in liquid nitrogen, and ground into a fine powder using an MM400 laboratory mill (Retsch, Germany) at freezing temperature and resuspended in 1 ml acetone. Pigments were extracted for 30 min with continuous shaking at 1000 rpm at 20 °C in the dark. The extract was centrifuged at 11500 *g* for 10 min at 4°C, and the supernatant was collected and passed through a PTFE 0.2 μm pore size syringe filter.

Quantification of carotenoids and Chl(a+b) was performed by HPLC using a Shimadzu Prominence HPLC system (Shimadzu, Kyoto, Japan) consisting of an LC-20AD pump, a DGU-20A degasser, a SIL-20AC automatic sample injector, CTO-20AC column thermostat, and a Nexera X2 SPD-M30A photodiode-array detector. Chromatographic separations were carried out on a Phenomenex Synergi 4 µm Hydro-RP 80Å, 250 x 4.6 mm column. 20 μl aliquots of acetonic extract was injected into the column and the pigments were eluted by a linear gradient from solvent A (acetonitrile, water, triethylamine, in a ratio of 9:1:0.01) to solvent B (ethyl acetate) followed by 15 min re-equlibration in solvent A. The gradient from solvent A to solvent B was run from 0 to 25 min at a flow rate of 1 ml/min. The column temperature was set to 25°C. Eluates were monitored in a wavelength range of 260 nm to 750 nm. Pigments were identified according to their retention time and absorption spectrum and quantified by integrated chromatographic peak area recorded at the wavelength of maximum absorbance for each kind of pigments using the corresponding molar decadic absorption coefficient (Jeffrey et al., 2005). The deepoxidation index of the xanthophyll cycle components was calculated as (zeaxanthin + ½ antheraxanthin)/(violaxanthin + anteraxanthin + zeaxanthin).

### Determination of non-photochemical quenching (NPQ) and the rate of electron donation from Asc to PSII

NPQ was determined using a Dual-PAM-100 instrument (Heinz Walz GmbH, Germany, Schreiber et al., 2008), on leaves dark-adapted for 30 min. In the initial seven minutes, the actinic light intensity was 100 µmol photons m^−2^ s^−1^, then the actinic light intensity was increased to 344 µmol photons m^−2^ s^−1^ for seven additional minutes. Saturating pulses of 5000 µmol photons m^−2^ s^−1^ were provided once every minute.

Fast Chl *a* fluorescence measurements were conducted at room temperature using a special version of the Handy-PEA instrument (Hansatech Instruments Ltd.) that allows for a reduction in measurement length to 5 ms. Leaf samples were heat-treated in a water bath (50°C, 40 s) to eliminate the oxygen-evolving activity, enabling the measurement of the rate of electron donation from Asc to PSII (Tóth et al., 2009). Following 15 min of dark adaptation, leaves were illuminated with continuous red light emitted by three LEDs (3500 µmol photons m^−2^ s^−1^, 650 nm peak wavelength). The first reliably measured point of the fluorescence transient, taken as F_0_, occurred at 20 µs. The measurement length was 5 ms, and the dark intervals between the first and second light pulses were 2.3, 9.6, 16.9, 31.5, 38.8, 53.4, 75.3, 100, 200, or 500 ms. The regeneration kinetics of the K-step, corresponding to the rate of electron transfer from Asc to PSII, were calculated as F_20µs_/F_300µs_, and the t_1/2_ values were plotted (as described in Tóth et al., 2009).

### Statistics for the physiological and molecular biology experiments

The presented data are based on at least three independent experiments. When applicable, averages and standard errors (±SE) were calculated. Statistical significance levels between the mutants and their background strains under each growth conditions were analyzed using Welch’s unpaired t-test. The significance levels are presented when applicable as * p<0.1.

### LC/MS analysis

For LC/MS analysis, leaf material was harvested from eight-week-old Col-0 and *pht4;4-3* mutant plants six hours into the photoperiod. Approximately 50-100 mg leaf material was collected, weighted and frozen in liquid nitrogen before grinding with a MM400 laboratory mill (Retsch, Germany) at 30 Hz for 1 min. After homogenisation, the samples were resuspended in extraction buffer (40 % (v/v) methanol, 20 % (v/v) water, 40 % (v/v) acetonitrile). After adding the extraction buffer (1 ml for 80 mg leaves), the samples were vortexed vigorously for 30 s. This was followed by centrifugation at 20000 *g*, at 4°C, for 30 min and the supernatant was collected. This step was repeated once.

For metabolome analysis, ultrahigh-performance liquid chromatography-tandem mass spectroscopy (UPLC-MS/MS) system, namely a Thermo Q-Exactive Focus instrument (Thermo Fisher Scientific, USA) equipped with a Dionex Unimate 3000 UHPLC system was used. For chromatographical separation an Xbidge Premier BEH Amide column (2.1 mm×150 mm, 2.5μm; Waters Corporation). Each sample was injected twice (2.5 µl), one for negative ionisation mode and one for the positive mode. For negative ionisation mode, the mobile phase was gradually changed from 99 % solvent A (97.5/2.5 acetonitrile / water to 80 % solvent B (20/80 acetonitrile

/ 20mM ammonium acetate + 20 mM ammonia in water) in 14 min using a flow of 400 μL/min. In positive ionisation mode, the mobile phase was gradually changed from 99 % solvent A (95/5 acetonitrile / 10mM Ammonium formate + 0.125% formic acid in water to 80 % solvent B (30/70 acetonitrile / 10mM Ammonium formate + 0.125% formic acid in water) in 11 min using a flow of 400 μL/min. For acquiring semi-quantitative metabolome data, system was operated in MS1 scan mode. Data were acquired using the following settings: sheath gas: 55 l/min, aux gas: 14 l/min, sweep gas: 4 l/min, probe heater 440°C, capillary temperature 280°C with 3500 V capillary voltage in positive ionisation mode and 2500V capillary voltage in negative ionisation mode. Scanning mass range was 100 to 900 m/z with an agc target 3*10^6 and maximum IT time 250 ms. Pooled extract samples as QCs were injected multiple time across the batch as suggested by the guidelines of Broadhurst et al. (2018).

For metabolite identification, pooled extract samles were injected multiple times at the end of the sequence. For acquiring MS2 data 10; 30; 50 NCE and 10; 20; 30 CE were used for DDMS and PRM modes, respectively, with automatic ion exclusion time.

### Data processing for metabolomics

To obtain peak intensity values, the recorded data were processed using MZmine2 (Puskal et al., 2010). Among MZmine modules, rolling ball baseline correction, FTMS peak filtration, ADAP chromatogram deconvolution, ADAP (wavelet) peak detection and peak alignment were used.

Metabolites are reported following recently updated recommendations (Alseekh et al., 2021). Briefly, MS1 level identification (analogous to level “D”) is based on exact mass matching between the given metabolite ion and the metabolites’s calculated monoisotopic mass from the *Arabidopsis thaliana* genome-scale metabolic network reconstruction (Dal’Molin et. al. 2010). Note that this procedure of metabolite identification is only putative as it may result in multiple equivalently probable metabolite assignments to the same peak, and the same identification may apply to multiple peaks. However, as this procedure is based on an organism-specific metabolic network reconstruction, it is suitable suitable to identify signals originating from *A. thaliana* among the 100s and 1000s mass spectrometric signals.

MS2 level identification (analogous with level “B” in Alseekh et al., 2021) is based on MS2 fragmentation spectra. For MS2 level metabolite identification Compound Discoverer 2.1 was used (similarly to e.g., Hemmer et al., 2020) with searching against the Endogenous Metabolites compound class of the mzCloud database (similarly to e.g., Klupczynska et al., 2017). Matching scores were accepted above 70 and hits were manually curated. Importantly, this procedure results in high confidence identifications that are only occasionally ambiguous. All metabolite identification data are reported in Suppl. File 7.

### Data analysis for metabolomics

All statistical procedures were performed using the R statistical language version 4.0. All data will be available in the Zenodo repository (https://zenodo.org/). Metabolite features were filtered based on detection frequency across multiple injected pooled extract samples (maximum non-detection frequency was set to 12.5%) and peak area reproducibility (coefficient of variation (CV) <20%). All data were normalized using Probabilistic Quotient Normalization (PQN, Dieterle et al., 2006). For PCA analysis, the missing values were input with the KNN method (DMwR2 package). PCA analysis was performed using the prcomp function of stats-package. Statistical analysis and plotting were performed using tidyverse (Wickham et al., 2019) and ggplot (Wickham et al., 2015). Size proportional Venn diagrams (Euler diagrams) were generated using the Euler function of the eulerr package (Larsson et al., 2021).

### Chloroplast extraction from Arabidopsis leaves for the PISA assay

Arabidopsis Col-0 plants were grown from seeds at 100 µmol photons s^−1^m^−2^ in 12 h day/night cycles for 5 weeks before harvest. Chloroplasts were isolated from leaves following Aronsson and Jarvis (2011) with some modifications. Leaves were cut and placed in ice-cold isolation buffer (0.3 M sorbitol, 5 mM MgCl_2_, 5 mM EGTA, 5 mM EDTA, 10 mM NaHCO_3_, 20 mM HEPES-KOH pH 8). Harvested leaves were homogenized with an IKA Ultra-Turrax T-25 disperser at 50 % power for 3 seconds, and then filtered through double-layered Miracloth. Retentate was resuspended in isolation buffer, homogenized, and filtered. Homogenization was repeated a total of 5 times. The filtered homogenate was centrifuged at 1000 g for 5 min at 4°C and the pellet was gently resuspended in isolation buffer. A Percoll gradient was prepared by centrifuging equal parts Percoll and isolation buffer with 0.6 mM glutathione at 43 000 g for 30 min at 4°C, which was subsequently used to separate intact and broken chloroplasts at 7800 g for 10 min at 4°C. Intact chloroplasts were recovered from the bottom layer, diluted in HMS buffer (50 mM HEPES-NaOH pH 8, 3 mM MgSO_4_, 0.3 M sorbitol) and pelleted at 1000 g for 5 min at 4°C. Chloroplasts were gently resuspended in 100 µL HMS buffer and illuminated with lamps at 400 µmol photons s^−1^m^−2^ for 5 min before being snap-frozen in liquid nitrogen. Chlorophyll content was measured in 80% acetone at 647 and 664 nm and chloroplast intactness was assessed through oxygen evolution as previously described (Joly and Carpentier 2011).

### Proteome integral solubility assay

Frozen chloroplast aliquots were thawed on ice and NP-40 Surfact-Amps™ Detergent Solution (ThermoFisher, cat. no 13444269) was added to 0.8 %. Chloroplasts were lysed with three cycles of bead beating (45 s, 6.5 m/s; FastPrep-24 5G) with cooling on ice between cycles. Lysates were cleared by centrifugation at (21,000 g, 5 min, 4 °C) and endogenous metabolites were removed with Zeba Spin Desalting Columns (ThermoFisher, cat. no 89882). The lysate was then split into sample replicates for PISA (four replicates for each treatment), with each replicate containing 200 µg of protein and diluted to a concentration of 1.25 µg/µL in lysis buffer (100 mM HEPES pH 8, 3 mM MgCl2, 150 mM KCl). Samples were then incubated in quadruplicates for 10 min under the following four conditions at protein concentration of 1 µg/µL: i) lysis buffer, ii) 2.5 mM Asc, ii) 5 mM Asc, and iv) 10 mM Asc. Concentrated solutions of Asc were adjusted to pH 8 in lysis buffer before addition to samples. Samples were then split and incubated in a thermocycler at 16 temperature points between 40-55 °C for 3 min. After allowing for precipitation at room temperature for 6 min, temperature points for each replicate were pooled and ultracentrifuged at 150 000 g for 30 min. The supernatants were then reduced with 8 mM DTT for 45 min followed by alkylation with 17 mM IAA for 30 min in darkness. Proteins were digested overnight (16 hours, 600 RPM) with Trypsin/Lys-C Protease Mix (Thermo Scientific, cat. No. A40009) at a 1:50 enzyme:protein ratio. Following digestion, samples were acidified to pH 2 with formic acid and desalted with pipette tips packed with six layers of C18 Empore™ SPE Disks (Merck, cat. no.66883-U) and dried in a Speed Vac at 45 °C for 1 hour. For TMT labeling of peptides, 12.5 µg of each sample was resuspended in 20 µL 20 mM EPPS buffer pH 8.5 and labeled with 0.1 mg TMTpro™ 16plex (Thermo Scientific, cat. no. A44521) for 1 hour at 600 rpm. Labeling was quenched by addition of hydroxylamine to 0.37 % for 15 min before pooling of samples. Labeled peptides were separated into 6 fractions with Pierce™ High pH Reversed-Phase Peptide Fractionation Kit (Thermofisher, Cat. no. 84868) and dried in a Speed Vac at 45 °C for 3.5 hours. Dried peptides were stored at −20 °C before LC-MS injection.

### LC/MS analysis of TMT labeled peptides

Peptides were reconstituted in 0.1% formic acid and analyzed on a Q-Exactive HF Hybrid Quadrupole-Orbitrap Mass Spectrometer connected to an UltiMate 3000 RSLCnano System with an EASY-Spray ion source. Peptides were loaded on a C18 Acclaim PepMap 100 trap column (75 μm x 2 cm, 3 μm, 100 Å) with a flow rate of 7 μL per min, using 3% acetonitrile, 0.1% formic acid and 96.9% water as solvent. Separation was performed using a ES802 EASY-Spray PepMap RSLC C18 Column (75 μm x 25 cm, 2 μm, 100 Å) at flow rate of 0.7 μL per minute and a 120 minutes linear gradient from 1% to 32% with 95% acetonitrile, 0.1% formic acid and 4.9% water as secondary solvent. Analysis was performed using one full scan (resolution 120,000 at 200 m/z, mass range 350-1500 m/z) followed by 15 MS2 DDA scans with the 15 most abundant peptides (resolution 60,000 at 200 m/z with an isolation window of 0.7 m/z). Precursor ions were fragmented with high-energy collision-induced dissociation at an NCE of 30. Maximum injection time and automatic gain control was set to 50 ms and 3E6 for MS1, and to 120 ms and 1E5 for MS2.

### Data analysis for the PISA assay

Protein identification and quantification was performed in MaxQuant version 2.2.0.0 using UniProt proteome UP000006548 as a library. Oxidation of methionine and N terminus acetylation was set as variable modifications and carbamidomethylation of cysteine was set as a fixed modification. A maximum of two missed cleavages were allowed and the false discovery rate was set to 1%. MSstats package version 4.4.1 (Choi, 2014) was used to median normalize proteins and calculate fold-changes in R version 4.2.2. p-values were adjusted for multiple hypothesis testing (Benjamini-Hochberg method) and a significance threshold of 0.05 was implemented. Proteins detected in less than 2 replicates were excluded. Quality assessment statistics and visualizations were generated using R version 4.2.2 and Tidyverse version 1.3.1 (Wickham et al., 2019). The complete dataset can be found in Suppl. File 6.

### Targeted quantification of Asc degradation intermediates by LC/MS

We followed an established method (Al Kadhi et al., 2017), with modifications for the quantification of Asc degradation intermediates. PISA samples were diluted 10-fold and filtered with Amicon® Ultra Centrifugal Filter, 3 kDa MWCO (Merck - UFC500324) and 0.5 µl were injected to LC-MS system. Asc degradation intermediates were separated using a Waters HSS T3 column (1.7 μm, 2.1 mm X 100 mm) on a liquid chromatography (Waters ACQUITY Premier) and tandem mass spectrometry (Waters TQS-Micro) system. Buffer A was composed of LC-MS grade water (VWR - 83645.320), 0.175% formic acid (Sigma 5.33002) and buffer B of 100 acetonitrile (VWR - 83640.320), The gradient elution was performed at a constant flow rate of 0.25 ml/min. Starting conditions were 100% A for 2.3 minutes. From 2.3 min to 3 min, B was increased to 30% with a gradient profile of 10, and kept for 0.5 min before returning to the initial conditions. The column was then equilibrated for another 1.2 minutes, resulting in a 4.7-minute chromatographic run. Compounds were identified by matching retention time (1.13 and 1.24 min for 2-keto-gulonic acid and tartatic acid, respectively) and fragmentation (transition: 193 > 58.9 and 149 > 87) with commercially available standards (89986846 – Molekula Group; T400-100g - Sigma-Aldrich). For MS/MS acquisition - 0.5kV capillary voltage, desolvation gas temperature was 600 °C and desolvation gas flow was 1000 l/min. Signals for organic acids were then acquired in MRM mode in Masslynx software. Quantification was performed using external calibration in the same sample matrix that was used for PISA assay (Rathod et al., 2020).

### Cloning of SBPase and thioredoxin F1

The SBPase gene (At3g55800) and the thioredoxin F1 gene (At3g02730) without transit peptides from *A. thaliana* were codon-optimized for *E. coli* and synthesized as geneblocks by Integrated DNA Technologies. The geneblocks were cloned into pET-28a(+) using Gibson assembly. After verification by sequencing the plasmids were transformed into *E. coli* BL21 using heat shock.

### Protein expression and purification

The BL21 mutants were cultivated in low salt LB media at 37 °C, 200 RPM until OD_600_ 0.4, whereafter overexpression was induced using 0.5 mM IPTG overnight at 30°C. Cells were harvested by centrifugation and stored at −20°C until further processing. Cell pellets were lysed in B-PERTM Complete Bacterial Protein Extraction Reagent (ThermoFischer Scientific) according to manufacturer instructions and cell debris was cleared by centrifugation. The supernatant was filtered through a 0.2 μm syringe filter before purification using an ÄKTA start protein purification system with a HisTrap Fast Flow Cytiva column (1 mL). The column was washed with 15 column volumes of wash buffer (50 mM Tris-HCl, 500 mM NaCl, 20 mM imidazole, pH 8) and eluted with a stepwise gradient of elution buffer (50 mM Tris-HCl, 500 mM NaCl, 300 mM imidazole, pH 8). Fractions containing protein were collected and the buffer was exchanged to storage buffer (50 mM Tris-HCl, pH 8) using a PD-10 Cytiva desalting column. Purified protein was stored at −80°C in aliquots.

### SBPase activity assay

A Malachite Green (MG) assay adapted from a previously published protocol (Vardakou et al., 2014) was used to assess the effect of Asc on purified SBPase enzyme. The assay measures the conversion rate of the alternative SBPase substrate fructose-1-6-bisphosphate to fructose-6-phosphate through the accumulation of inorganic phosphate. Phosphate colometric development solution was prepared fresh by mixing 400 µL MG dye stock (1.55 g/L Malachite Green oxalate salt, 3 M H2SO4) with 125 µL ammonium molybdate (60 mM) and 10 µL Tween-20 (11% v/v)).

The solution was filtered through a 0.2 µm filter and kept dark. Development plates were prepared by mixing 36 µL development solution with 100 µL reaction buffer (50 mM Tris-Hcl, 15 mM MgCl2, 10 mM DTT) in 96-well plates and kept dark. Frozen aliqots of SBPase and thioredoxin F1 was thawed on ice and centrifuged at 25,000 xg for 5 minutes to remove precipitated protein. SBPase enzyme was activated for 1 hour at room temperature by incubation in reaction buffer with 20 µM thioredoxin F1. To measure the effect of Asc, 25 µL activated SBPase solution containing 8.4 ng/µL SBPase was incubated for 15 minutes with 25 µL reaction buffer with and without 20 µM Asc in quadruplicates. The reaction was initiated by multipipetting 50 µL fructose-1-6-bisphosphate to a final concentration of 500 µM and 20 µL of the mixture was immediatly sampled to the prepared development plate. Triplicate phosphate standards of 0, 25, 50 and 100 µM was pipetted (20 µL) to the development plate. To stabilize the color, 7.5 µL sodium citrate (34% w/v) was added to the development plate and the absorbance was measured at 620 nm after 20 minutes. The sampling of the enzyme reaction mixture was repeated after 30 minutes with fresh phosphate standards.

## Author contributions

S.Z.T., B.P. and E.P.H designed the research. D.T., R.T., F.A., A.K., A. V.-M., L.K., and S.K. performed research. D.T., R.T., A.K., L.K., A.R.F., E.P.H., B.P. and S.Z.T. analysed data. S.Z.T. with the contributions of D.T., R.T., A.K., A.R.F, E.P.H. and B.P. wrote the article.

## Acknowledgements

The authors are grateful to Fredrik Edfors (KTH, Solna, Sweden) for assistance with LC/MS for the PISA assay, and Csilla Sajben (HUN-REN BRC, Szeged) for the assistance with the metabolomics analyses.

## Data availability

Raw metabolite profiling data and the complete dataset of the PISA assay are provided in Suppl. Files 6 and 7.

## Supplementary Materials

**Suppl. Fig. 1.** Characterization of *PHT4;4* and *VTC2* mutants of *A. thaliana*. **A)** RT-PCR performed using primers annealing downstream of the predicted insertion site in *PHT4;4* (primers P7 and P9), and *VTC2*. The expected sizes are marked with arrows. Ubiquitin was used as a reference gene, Ctrl stands for water-control. The number of RT-PCR cycles was 30. **B)** Phenotypes of eight-week-old *pht4;4-1*, *pht4;4-3, pht4;4-4,* and *vtc2-4* mutants and their respective WTs (Ler-0 and Col-0), grown at NL. **C)** Rosette sizes of eight-week-old plants grown at NL. The averages are based on six independent experiments, with 20 to 25 replicates in each. B). Statistical significance levels between the mutants and their background strains were analyzed using Welch’s unpaired t-test. The significance levels are presented as *. Primers are listed in Suppl. Table 1.

**Suppl. Fig. 2.** Assessment of chloroplastic Asc content by three independent approaches. A) Boxplot representation of the total Asc content, distributed among the chloroplast, the cytosol and the vacuole, in the *vtc2-4* and *pht4;4-3* mutants and Col-0, assessed by non-aqueous fractionation. B) Asc content of chloroplasts isolated from the *vtc2-4* and *pht4;4-3* mutants and Col-0, quantified by HPLC. The chloroplast intactness was in the range of 80% (see Materials and Methods). C) The half-time (t_1/2_) of electron transfer from Asc to PSII in heat-treated (50°C, 40 s) leaves of *vtc2-4, vtc2-1,* and *pht4;4-3* mutants and Col-0. The t_1/2_ values were derived from the regeneration kinetics of the K step of the fast chl *a* fluorescence transient (calculated as F_20μs_/F_300μs_; see Tóth et al., 2009). Data represent averages of 16 independent biological replicates with standard errors. The significance of differences between means were determined by ANOVA with Tukey post-hoc test. Different letters indicate significant differences between means (P < 0.1).

**Suppl. Fig. 3.** Principal component analysis performed on Col-0, *vtc2-4*, and *pht4;4-3* mutant lines. The analysis was performed based on 244 metabolites, putatively annotated at MS1 and/or MS2 levels (levels “B” and “D”, see Materials and Methods and Suppl. File 3).

**Suppl. Fig. 4.** Concentration of Asc and its degradation products in PISA-samples to which 10 mM Asc was added, and then incubated at room temperature (RT) or heat-treated at 55°C for 3 min (HT). A) HPLC analysis of Asc and DHA concentrations in the treated chloroplasts B) Concentration of the degradation products of Asc (2-keto-gulonic acid and L-tartaric acid), detected by mass spectrometry.

**Suppl. Text 1.** Detailed description of metabolite changes in the *vtc2-4* and *pht4;4-3* mutants and Col-0.

**Suppl. Text 2.** Additional putative interactions between Asc and chloroplastic proteins indicated by the PISA assay.

**Suppl. Table 1.** List of primer pairs used in this study.

**Suppl. Table 2.** Metabolite changes specific to the *vtc2-4* mutant.

**Suppl. File 1**

Supplementary file 1 contains the detailed results of RT-qPCR analysis, carotenoids and Asc contents and Chl(a+b) content determinations.

**Suppl. File 2**

Supplementary file 2 contains metabolite identifications and corresponding peak identifiers, (row_ID), retention time, peak average mass / charge ratio (mzMed), theoretical mass / charge ratio (theoretical_mz), mass measurement error (mass_difference_ppm) and metabolite data. If the identification is based on only exact mass (level “D”) and multiple alternative putative identifications are possible, corresponding Kegg IDs and alternative identifications are listed and separated with “|”.

**Suppl. File 3**

Supplementary file 3 contains metabolite intensity fold-change differences and p-values (from two sided t-tests) before and after false discovery rate correction between all pairs of Col-0, *vtc2-4* and *pht4;4-3* lines. Metabolites were identified based on exact mass data (level “D”) and MS2 fragmentation pattern data (level “B”), see Materials and Methods.

**Suppl. File 4**

Supplementary file 4 contains metabolite intensity fold-change differences and p-values (from two sided t-tests) before and after false discovery rate correction between Col-0 / *vtc2-4* and Col-0 / *pht4;4-3* pairs. The metabolites were identified based on their MS2 fragmentation pattern data (level “B”); see Materials and Methods.

**Suppl. File 5**

Supplementary file 5 contains information on the PISA assay: List of the significant proteins at 2, 5 and 10 mM Asc along with log2FC, standard error (SE), p-values, adjusted p-values (Benjamini-Hochberg method), and Uniprot annotations.

**Suppl. File 6**

Supplementary file 6 contains the complete dataset on the PISA assay at 2, 5 and 10 mM Asc. It contains log2FC, standard error (SE), p-values, adjusted p-values (Benjamini-Hochberg method), and Uniprot annotations for detected proteins.

**Suppl. File 7**

Supplementary File 7 contains metabolite ion intensity data (intensity) with corresponding peak identifier (raw_ID) and identifications – (MS1_metabolite_name (level “D”); MS2_metabolite_name (level “B”)), according to the sample groups (class), biological replicates (Sample_ID) and measurement raw MS files (sample).

## References

Agius F, Gonzalez-Lamothe R, Caballero JL, Munoz-Blanco J, Botella MA, Valpuesta V (2003) Engineering increased vitamin C levels in plants by overexpression of a D-galacturonic acid reductase. Nature Biotechnol 21: 177–181

Al Kadhi O, Melchini A, Mithen R, Saha S (2017) Development of a LC-MS/MS method for the simultaneous detection of tricarboxylic acid cycle intermediates in a range of biological matrices. J Anal Methods Chem 2017: 5391832

Alseekh S, Aharoni A, Brotman Y, Contrepois K, D’Auria J, Ewald J, C Ewald J, Fraser PD, Giavalisco P, Hall RD, Heinemann M, Link H, Luo J, Neumann S, Nielsen J, Perez de Souza L, Saito K, Sauer U, Schroeder FC, Schuster S, Siuzdak G, Skirycz A, Sumner LW, Snyder MP, Tang H, Tohge T, Wang Y, Wen W, Wu S, Xu G, Zamboni N, Fernie AR (2021) Mass spectrometry-based metabolomics: a guide for annotation, quantification and best reporting practices. Nat Methods 18: 747–756

Alvarez ME, Savouré A, Szabados L (2021) Proline metabolism as regulatory hub. Trends Plant Sci 27: 39–55

Araújo WL, Martins AO, Fernie AR, Tohge T (2014) 2-oxoglutarate: linking TCA cycle function with amino acid, glucosinolate, flavonoid, alkaloid, and gibberellin biosynthesis. Front Plant Sci 5: 552

Aronsson H, Jarvis RP (2011) Rapid isolation of Arabidopsis chloroplasts and their use for in vitro protein import assays. In: Jarvis RP (ed) Chloroplast Research in Arabidopsis: Methods and Protocols, Volume I. Humana Press, Totowa, NJ, pp 281–305

Asada K (2006) Production and scavenging of reactive oxygen species in chloroplasts and their functions. Plant Physiol 141: 391–396

Ashihara H, Ludwig IA, Katahira, R, Yokota T, Fujimura T, Crozier A (2015) Trigonelline and related nicotinic acid metabolites: occurrence, biosynthesis, taxonomic considerations, and their roles in planta and in human health. Phytochem Rev 14: 765–798

Barth C, De Tullio M, Conklin PL (2006). The role of ascorbic acid in the control of flowering time and the onset of senescence. J Exp Bot 57: 1657–1665

Bartoli CG, Tambussi EA, Diego F, Foyer CH (2009) Control of ascorbic acid synthesis and accumulation and glutathione by the incident light red/far red ratio in *Phaseolus vulgaris* leaves. FEBS Lett 583: 118–122

Baxter CJ, Redestig H, Schauer N, Repsilber D, Patil KR, Nielsen J, Selbig J, Liu J, Fernie AR, Sweetlove LJ (2007) The metabolic response of heterotrophic Arabidopsis cells to oxidative stress. Plant Physiol 143: 312–325

Bouchnak I, Brugière S, Moyet L, Le Gall S, Salvi D, Kuntz M, Tardif M, Rolland N (2019) Unraveling hidden components of the chloroplast envelope proteome: Opportunities and limits of better MS sensitivity. Mol Cell Proteomics 18: 1285–1306

Bratt C, Arvidsson P, Carlsson M, Akerlund H (1995) Regulation of violaxanthin de-epoxidase activity by pH and ascorbate. Photosynth Res 45: 169–175

Broadhurst, D., Goodacre, R., Reinke, S.N. (2018) Guidelines and considerations for the use of system suitability and quality control samples in mass spectrometry assays applied in untargeted clinical metabolomic studies. Metabolomics 14:72

Bulley S, and Laing W (2016) The regulation of ascorbate biosynthesis. Curr Opin Plant Biol 33: 15–22

Chen H, Xiong L (2005) Pyridoxine is required for post-embryonic root developmentand tolerance to osmotic and oxidative stresses. Plant J 44: 396–408

Chen Y, Hoehenwarter W (2015) Changes in the phosphoproteome and metabolome link early signaling events to rearrangement of photosynthesis and central metabolism in salinity and oxidative stress response in Arabidopsis. Plant Physiol 169: 3021–3033

Chen Z, Gallie DR (2004) The ascorbic acid redox state controls guard cell signaling and stomatal movement. Plant Cell 16: 1143–1162

Chen, M, Thelen JJ (2011) Plastid uridine salvage activity is required for photoassimilate allocation and partitioning in Arabidopsis. Plant Cell 23: 2991–3006

Che-Othman MH, Jacoby RP, Millar AH, Taylor NL (2020) Wheat mitochondrial respiration shifts from the tricarboxylic acid cycle to the GABA shunt under salt stress. New Phytol 225: 1166–1180

Conklin PL, Saracco SA, Norris SR, Last RL (2000) Identification of ascorbic acid-deficient *Arabidopsis thaliana* mutants. Genetics 154: 847–856

Crosatti C, Rizza F, Badeck FW, Mazzucotelli E, Cattivelli L (2013) Harden the chloroplast to protect the plant. Physiol Plantarum 147: 55–63

Dastogeer KMG, Li H, Sivasithamparam K, Jones M, Du X, Ren Y, Wylie SJ (2017) Metabolic responses of endophytic *Nicotiana benthamiana* plants experiencing water stress. Environ Exp Bot 143: 59–71

de Oliveira Dal’Molin CG, Quek LE, Palfreyman RW, Brumbley SM, Nielsen LK (2010) AraGEM, a genome-scale reconstruction of the primary metabolic network in Arabidopsis. Plant Physiol 152: 579–589

Dieterle F, Ross A, Schlotterbeck G, Senn H (2006) Probabilistic Quotient Normalization as robust method to account for dilution of complex biological mixtures. Application in ^1^H NMR metabonomics. Anal Chem 78: 4281–4290

Ding F, Wang M, Zhang S (2017) Overexpression of a Calvin cycle enzyme SBPase improves tolerance to chilling-induced oxidative stress in tomato plants. Sci Hortic 214: 27–33

Dowdle J, Ishikawa T, Gatzek S, Rolinski S, Smirnoff N (2007) Two genes in *Arabidopsis thaliana* encoding GDP-L-galactose phosphorylase are required for ascorbate biosynthesis and seedling viability. Plant J 52: 673–689

Draper J, J. Lloyd A, Goodacre R, Beckmann M (2013) Flow infusion electrospray ionisation mass spectrometry for high throughput, non-targeted metabolite fingerprinting: a review. Metabolomics 9: 4–29

Farrow SC, Facchini PJ (2004) Functional diversity of 2-oxoglutarate/Fe(II)-dependent dioxygenases in plant metabolism. Front Plant Sci 5: 524

Feldman-Salit A, Veith N, Wirtz M, Hell R, Kummer U (2019) Distribution of control in the sulfur assimilation in *Arabidopsis thaliana* depends on environmental conditions. New Phytol 222: 1392–1404

Feller, U (2016) Drought stress and carbon assimilation in a warming climate: Reversible and irreversible impacts. J Plant Physiol 203: 69–79

Fernie AR, Tóth SZ (2015) Identification of the elusive chloroplast ascorbate transporter extends the substrate specificity of the PHT family. Mol Plant 8: 674–676

Fotopoulos V, Sanmartin M, Kanellis AK (2006). Effect of ascorbate oxidase over-expression on ascorbate recycling gene expression in response to agents imposing oxidative stress. J Exp Bot 57: 3933–3943

Foyer CH, Kyndt T, Hancock RD (2020) Vitamin C in plants: Novel concepts, new perspectives, and outstanding issues. Antioxid Redox Signal 32: 463–485

Foyer CH, Lelandais MA (1996) A comparison of the relative rates of transport of ascorbate and glucose across the thylakoid, chloroplast and plasmalemma membranes of pea leaf mesophyll cells. J Plant Physiol 148: 391–398

Gaetani M, Sabatier P, Saei AA, Beusch CM, Yang Z, Lundstrom SL, Zubarev RA (2019) Proteome integral solubility alteration: a high-throughput proteomics assay for target deconvolution. J Proteome Res 18: 4027–4037

Gakière B, Hao J, de Bont L, Pétriacq P, Nunes-Nesi A, Fernie AR (2018) NAD^+^ biosynthesis and signaling in plants. Crit Rev Plant Sci 37: 259–307

Gjindali A, Johnson GN (2023) Photosynthetic acclimation to changing environments. Biochem Soc Trans 51: 473–486

Graciet E, Lebreton S, Gontero B (2004) Emergence of new regulatory mechanisms in the Benson-Calvin pathway via protein-protein interactions: a glyceraldehyde-3-phosphate dehydrogenase/CP12/phosphoribulokinase complex. J Exp Bot 55: 1245–1254

Guo B, Jin Y, Wussler C, Blancaflor EB, Motes CM, Versaw WK (2008) Functional analysis of the Arabidopsis PHT4 family of intracellular phosphate transporters. New Phytol 177: 889–898

Gururani MA, Venkatesh J, Tran LSP (2015) Regulation of photosynthesis during abiotic stress-induced photoinhibition. Mol Plant 8: 1304–1320

Gütle DD, Roret T, Müller SJ, Couturier J, Lemaire SD, Hecker A, Dhalleine T, Buchanan BB, Reski R, Einsle O, Jacquot JP (2016) Chloroplast FBPase and SBPase are thioredoxin-linked enzymes with similar architecture but different evolutionary histories. Proc Nat Acad Sci USA 113: 6779–6784

Hallin EI, Guo K, Åkerlund H-E (2016) Functional and structural characterization of domain truncated violaxanthin de-epoxidase. Physiol Plantarum 157: 414–421

Havaux M, Ksas B, Szewczyk A, Rumeau D, Franck F, Caffarri S, Triantaphylidès C (2009) Vitamin B6 deficient plants display increased sensitivity to high light and photo-oxidative stress. BMC Plant Biol 9: 130

Hemmer S, Manier SK, Fischmann S, Westphal F, Wagmann L, Meyer MR (2020) Comparison of three untargeted data processing workflows for evaluating LC-HRMS metabolomics data. Metabolites 10: 378

Hildebrandt TM (2018) Synthesis versus degradation: Directions of amino acid metabolism during Arabidopsis abiotic stress response. Plant Mol Biol 98: 121–135

Hoang MTT, Almeida D, Chay S, Alcon C, Corratge-Faillie C, Curie C, Mari S (2021) AtDTX25, a member of the multidrug and toxic compound extrusion family, is a vacuolar ascorbate transporter that controls intracellular iron cycling in Arabidopsis. New Phytol 231: 1956–1967

Irigoyen S, Karlsson PM, Kuruvilla J, Spetea C, Versaw WK (2011) The sink-specific plastidic phosphate transporter PHT4;2 influences starch accumulation and leaf size in Arabidopsis. Plant Physiol 157: 1765–1777

Ishikawa T, Takahara K, Hirabayashi T, Matsumura H, Fujisawa S, Terauchi R, Uchimiya H, Kawai-Yamada M (2009) Metabolome analysis of response to oxidative stress in rice suspension cells overexpressing cell death suppressor Bax inhibitor-1. Plant Cell Physiol 51: 9–20

Ivanov B, Asada K, Edwards GE (2007) Analysis of donors of electrons to photosystem I and cyclic electron flow by redox kinetics of P700 in chloroplasts of isolated bundle sheath strands of maize. Photosynth Res 92: 65–74

Ivanov BN, Sacksteder CA, Kramer DM, Edwards GE (2001) Light-induced ascorbate-dependent electron transport and membrane energization in chloroplasts of bundle sheath cells of the C_4_ plant maize. Arch Biochem Biophys 385: 145–153

Jamai A, Salomé PA, Schilling SH, Weber APM, McClung CR (2009) Arabidopsis photorespiratory serine hydroxymethyltransferase activity requires the mitochondrial accumulation of Ferredoxin-Dependent Glutamate Synthase. Plant Cell 21: 595–606

Jeffrey SW, Mantoura RFC, Wright SW (1997) Phytoplankton pigments in oceanography: guidelines to modern methods. UNESCO Publishing, Paris

Kang Z, Qin T, Zhao Z (2019) Thioredoxins and thioredoxin reductase in chloroplasts: A review. Gene 706: 32–42

Kavkova I, Blöchl C, Tenhaken R (2019) The Myo-inositol pathway does not contribute to ascorbic acid synthesis. Plant Biol 21: 95–102

Khan MS, Haas FH, Samami AA, Gholami AM, Bauer A, Fellenberg K, Reichelt M, Hänsch R, Mendel RR, Meyer AJ, Wirtz M, Hell R (2010) Sulfite reductase defines a newly discovered bottleneck for assimilatory sulfate reduction and is essential for growth and development in *Arabidopsis thaliana*. Plant Cell 22: 1216–1231

Kim HK, Verpoorte R (2010) Sample preparation for plant metabolomics. Phytochem Anal 21: 4–13

Kleine T, Nägele T, Neuhaus HE, Schmitz-Linneweber C, Fernie AR, Geigenberger P, Grimm B, Kaufmann K, Klipp E, Meurer J, Möhlmann T, Mühlhaus T, Naranjo B, Nickelsen J, Richter A, Ruwe H, Schroda M, Schwenkert S, Trentmann O, Willmund F, Zoschke R, Leister D (2021) Acclimation in plants – the Green Hub consortium. Plant J 106: 23–40

Klie D, Krueger S, Krall L, Giavalisco P, Flügge U-I, Willmitzer L, Steinhauser D (2011) Analysis of the compartmentalized metabolome – a validation of the non-aqueous fractionation technique. Front Plant Sci 2: 55

Klupczynska A, Dereziński P, Garrett TJ, Rubio VY, Dyszkiewicz W, Kasprzyk M, Kokot ZJ (2017) Study of early stage non-small-cell lung cancer using Orbitrap-based global serum metabolomics. J Cancer Res Clin Oncol 143, 649–659

Kovács L, Vidal-Meireles A, Nagy V, Tóth SZ (2016) Quantitative determination of ascorbate from the green alga Chlamydomonas reinhardtii by HPLC. Bio-Protoc 6:e2067

Krueger S, Giavalisco P, Krall L, Steinhauser M-C, Büssis D, Usadel B, Flügge U-I, Fernie AR, Willmitzer L, Steinhauser D (2011) A topological map of the compartmentalized *Arabidopsis thaliana* leaf metabolome. PLoS ONE 6: e17806

Krueger S, Giavalisco P, Krall L, Steinhauser M-C, Büssis D, Usadel B, Flügge U-I, Fernie A-R, Willmitzer L, Steinhauser D (2011) A Topological map of the compartmentalized *Arabidopsis thaliana* leaf metabolome. PLoS ONE 6: e17806

Krueger S, Steinhauser D, Lisec J, Giavalisco P (2014) Analysis of subcellular metabolite distributions within *Arabidopsis thaliana* leaf tissue: a primer for subcellular metabolomics. Methods Mol Biol 1062: 575–596

Larsson J, Godfrey AJR, Gustafsson P, Eberly DH, Huber E, Slowikowski K, Privé F (2021) eulerr: Area-proportional Euler and Venn diagrams with ellipses. https://cran.r-project.org/web/packages/eulerr/

Lehmann M, Schwarzländer M, Obata T, Sirikantaramas S, Burow M, Olsen CE, Tohge T, Fricker MD, Møller BL, Fernie AR, Sweetlove LJ, Laxa M (2009) The metabolic response of Arabidopsis roots to oxidative stress is distinct from that of heterotrophic cells in culture and highlights a complex relationship between the levels of transcripts, metabolites, and flux. Mol Plant 2: 390–406

Joly D, Carpentier R (2011) Rapid isolation of intact chloroplasts from spinach leaves. In: Carpentier R (ed) Photosynthesis Research Protocols. Methods in Molecular Biology, vol 684. Humana Press, Totowa, NJ. 10.1007/978-1-60761-925-3_24

Leister D (2019) Piecing the puzzle together: The central role of reactive oxygen species and redox hubs in chloroplast retrograde signaling. Antiox Redox Signal 30: 1206–1219

Li S, Park Y, Duraisingham S, Strobel FH, Khan N, Soltow QA, Jones DP, Pulendran B (2013) Predicting network activity from high throughput metabolomics. PLoS Comput Biol 9: e1003123

Liu X-L, Yu H-D, Guan Y, Li J-K, Guo F-Q (2012) Carbonylation and loss-of-function analyses of SBPase reveal its metabolic interface role in oxidative stress, carbon assimilation, and multiple aspects of growth and development in Arabidopsis. Mol Plant 5: 1082–1099

Li Z, Peers G, Dent RM, Bai Y, Yang SY, Apel W, Leonelli L, Niyogi KK (2016) Evolution of an atypical de-epoxidase for photoprotection in the green lineage. Nat Plants 2: 16140

Lim B, Smirnoff N, Cobbett CS, Golz JF (2016) Ascorbate-deficient *vtc2* mutants in Arabidopsis do not exhibit decreased growth. Front Plant Sci 7:1025

Linster CL, Adler LN, Webb K, Christensen KC, Brenner C, Clarke SG (2008) A second GDP-L-galactose phosphorylase in *Arabidopsis* en route to vitamin C - Covalent intermediate and substrate requirements for the conserved reaction. J Biol Chem 27: 18483–18492

Lorence A, Chevone BI, Mendes P, Nessler CL (2004) myo-Inositol oxygenase offers a possible entry point into plant ascorbate biosynthesis. Plant Physiol 134: 1200–1205

Malone LA, Proctor MS, Hitchcock A, Hunter CN, Johnson MP (2021) Cytochrome b_6_f – Orchestrator of photosynthetic electron transfer. Biochim Biophys Acta - Bioenergetics 1862: 148380

Malone LA, Qian P, Mayneord GE, Hitchcock A, Farmer DA, Thompson RF, Swainsbury DJK, Ranson NA, Hunter CN, Johnson MP (2019) Cryo-EM structure of the spinach cytochrome b_6_f complex at 3.6 Å resolution. Nature 575: 535–539

Mano J, Hideg É, Asada K (2004) Ascorbate in thylakoid lumen functions as an alternative electron donor to photosystem II and photosystem I. Arch Biochem Biophys 429: 71–80

Marri K, Zaffagnini M, Collin V, Issakidis-Bourguet E, Lemaire SD, Pupillo P, Sparla F, Miginiac-Maslow M, Trosta P (2009) Prompt and easy activation by specific thioredoxins of Calvin cycle enzymes of *Arabidopsis thaliana* associated in the GAPDH/CP12/PRK supramolecular complex. Mol Plant 2: 259–269

Mateus A, Kurzawa N, Becher I, Sridharan S, Helm D, Stein F, Typas A, Savitski MM (2020) Thermal proteome profiling for interrogating protein interactions. Mol Syst Biol 16: e9232

Medeiros DB, Arrivault S, Alpers J, Fernie AR, Aarabi F (2019) Non-aqueous (NAF) for metabolite analysis in subcellular compartments of Arabidopsis leaf tissues. Bio-Protoc 9: e3399

Miyaji T, Kuromori T, Takeuchi Y, Yamaji N, Yokosho K, Shimazawa A, Sugimoto E, Omote H, Ma JF, Shinozaki K, Moriyama Y (2015) AtPHT4;4 is a chloroplast-localized ascorbate transporter in Arabidopsis. Nature Comm 6: 5928

Müller-Moulé P, Conklin PL, Niyogi KK (2002) Ascorbate deficiency can limit violaxanthin de-epoxidase activity in vivo. Plant Physiol 128: 970–977

Müller-Moulé P, Golan T, Niyogi KK (2004) Ascorbate-deficient mutants of Arabidopsis grow in high light despite chronic photooxidative stress. Plant Physiol 134: 1163–1172

Müller-Moulé P, Havaux M, Niyogi KK (2003) Zeaxanthin deficiency enhances the high light sensitivity of an ascorbate-deficient mutant of Arabidopsis. Plant Physiol 133: 748–760

Murphy JT, Bruinsma JJ, Schneider DL, Collier S, Guthrie J, Chinwalla A, Robertson JD, Mardis ER, Kornfeld K (2011) Histidine protects against zinc and nickel toxicity in *Caenorhabditis elegans*. PLoS Genet 7: e1002013

Nazar R, Umar S, Khan NA (2015) Exogenous salicylic acid improves photosynthesis and growth through increase in ascorbate-glutathione metabolism and S assimilation in mustard under salt stress. Plant Signal Behav 10: e1003751.

Neubauer C, Yamamoto HY (1994) Membrane barriers and Mehler-peroxidase reaction limit the ascorbate available for violaxanthin de-epoxidase activity in intact chloroplasts. Photosynth Res 39: 137–147

Noctor G, Reichheld J-P, Foyer CH (2018) ROS-related redox regulation and signaling in plants. Semin Cell Dev Biol 80: 3–12

Parra M, Stahl S, Hellmann H (2018) Vitamin B6 and its role in cell metabolism and physiology. Cells 7: 84

Patel J, Ariyaratne M, Ahmed S, Ge L, Phuntumart V, Kalinoski A, Morris PF (2017) Dual functioning of plant arginases provides a third route for putrescine synthesis. Plant Sci 262: 62–73

Pluskal T, Castillo S, Villar-Briones A, Orešič M (2010) MZmine 2: modular framework for processing, visualizing, and analyzing mass spectrometry-based molecular profile data. BMC Bioinform 11: 1–11

Podmaniczki A, Nagy V, Vidal-Meireles A, Tóth D, Patai R, Kovács L, Tóth SZ (2021) Ascorbate inactivates the oxygen-evolving complex in prolonged darkness. Physiol Plantarum 171: 232–245

Rathod R, Gajera B, Nazir K, Wallenius J, Velagapudi V (2017) Simultaneous measurement of tricarboxylic acid cycle intermediates in different biological matrices using liquid chromatography–tandem mass spectrometry; Quantitation and comparison of TCA cycle intermediates in human serum, plasma, Kasumi-1 cell and murine liver tissue. Metabolites 10: 103

Porra RJ, Thompson WA, Kriedeman PE (1989) Determination of accurate extinction coefficients and simultaneous equations for essaying chlorophylls-a and -b with four different solvents: verification of the concentration of chlorophyll standards by atomic absorption spectroscopy. Biochim Biophys Acta 975: 384–394

Riedel A, Rutherford AW, Hauskall G, Muller A, Nitschke W (1991) Chloroplast Rieske Center. EPR study on its spectral characteristics, relaxation and orientation properties. J Biol Chem 266: 17838–17844

Rosado-Souza L, Fernie AR, Aarabi F (2020) Ascorbate and thiamin: Metabolic modulators in plant acclimation responses. Plants 9: 101

Ruban AV (2016) Nonphotochemical chlorophyll fluorescence quenching: mechanism and effectiveness in protecting plants from photodamage. Plant Physiol 170: 1903–1916

Ruiz-Pavón L, Lundh F, Lundin B, Mishra A, Persson BL, Spetea C (2008) Arabidopsis ANTR1 is a thylakoid Na^+^-dependent phosphate transporter: functional characterization in *Escherichia coli*. J Biol Chem 283: 13520–13527

Saga G, Giorgetti A, Fufezan C, Giacometti GM, Bassi R, Morosinotto T (2010) Mutation analysis of violaxanthin de-epoxidase identifies substrate-binding sites and residues involved in catalysis. J Biol Chem 285: 23763–23770

Savchenko T, Tikhonov K (2021) Oxidative stress-induced alteration of plant central metabolism. Life 11: 304

Savitski MM, Reinhard FB, Franken H, Werner T, Savitski MF, Eberhard D, Martinez Molina D, Jafari R, Dovega RB, Klaeger S, Kuster B, Nordlund P, Bantscheff M, Drewes G (2014) Tracking cancer drugs in living cells by thermal profiling of the proteome. Science 346: 1255784

Schansker G, Tóth SZ, Holzwarth AR, Garab G (2014) Chlorophyll *a* fluorescence: beyond the limits of the Q_A_ model. Photosynth Res 120: 43–58

Schippers JH, Nunes-Nesi A, Apetrei R, Hille J, Fernie AR, Dijkwel PP (2008) The *Arabidopsis onset of leaf death5* mutation of quinolinate synthase affects nicotinamide adenine dinucleotide biosynthesis and causes early ageing. Plant Cell 20: 2909–2925

Schöttler MA, Tóth SZ (2014) Photosynthetic complex stoichiometry dynamics in higher plants: environmental acclimation and photosynthetic flux control. Front Plant Sci 5: 188

Schreiber U, Klughammer C (2008) Non-photochemical fluorescence quenching and quantum yields in PSI and PSII: Analysis of heat-induced limitations using Maxi-Imaging-PAM and Dual-PAM-100. PAM Appl Notes 1: 15–18

Schwenkert S, Fernie AR, Geigenberger P, Leister D, Möhlmann T, Naranjo B, Neuhaus HE (2022) Chloroplasts are key players to cope with light and temperature stress. Trends Plant Sci. 27: 577–587

Shapiguzov A, Vainonen JP, Hunter K, Tossavainen H, Tiwari A, Järvi S, Hellman M, Aarabi F, Alseekh S, Wybouw B, Van Der Kelen K, Nikkanen L, Krasensky-Wrzaczek J, Sipari N, Keinänen M, Tyystjärvi E, Rintamäki E, De Rybel B, Salojärvi J, Van Breusegem F, Fernie AR, Brosché M, Permi P, Aro E-M, Wrzaczek M, Kangasjärvi J (2019) Arabidopsis RCD1 coordinates chloroplast and mitochondrial functions through interaction with ANAC transcription factors. eLife 8: e43284

Siddappa S, Marathe GK (2020) What we know about plant arginases? Plant Physiol Biochem. 156: 600–610

Sipari N, Lihavainen J, Shapiguzov A, Kangasjärvi J, Keinänen M (2020) Primary metabolite responses to oxidative stress in early-senescing and paraquat resistant *Arabidopsis thaliana rcd1* (*Radical-Induced Cell Death1*). Front Plant Sci 11: 194

Slocum RD (2005) Genes, enzymes and regulation of arginine biosynthesis in plants. Plant Physiol Biochem 43: 729–745

Smirnoff N (2018) Ascorbic acid metabolism and functions: A comparison of plants and mammals. Free Radic Biol Med 122: 116–129

Smirnoff N, Wheeler GL (2024) The ascorbate biosynthesis pathway in plants is known, but there is a way to go with understanding control and functions. J Exp Bot erad505

Sporre E, Karlsen J, Schriever K, Samuelsson JA, Janasch M, Kotol D, Strandberg L, Zeckey L, Piazza I, Syrén P-O, Edfors F, Hudson EP (2022) Metabolite interactions in the bacterial Calvin cycle and implications for flux regulation. bioRxiv 2022.03.15.483797

Sridharan S, Kurzawa N, Werner T, Günthner I, Helm D, Huber W, Bantscheff M, Savitski MM (2019) Proteome-wide solubility and thermal stability profiling reveals distinct regulatory roles for ATP. Nat Commun 10: 1155

Stirbet A, Lazár D, Kromdijk J, Govindjee (2018) Chlorophyll *a* fluorescence induction: Can just a one-second measurement be used to quantify abiotic stress responses? Photosynthetica 56: 86–104

Suss KH, Arkona C, Manteuffel R, Adler K (1993) Calvin cycle multienzyme complexes are bound to chloroplast thylakoid membranes of higher plants in situ. Proc Nat Acad Sci USA 90: 5514–5518

Telman W, Dietz K-J (2019) Thiol redox-regulation for efficient adjustment of sulfur metabolism in acclimation to abiotic stress. J Exp Bot 70: 4223–4236

Thakur M, Anand A (2021) Hydrogen sulfide: An emerging signaling molecule regulating drought stress response in plants. Physiol Plant 172: 1227–1243

Tóth SZ (2023) The functions of chloroplastic ascorbate in vascular plants and algae. Int J Mol Sci 24: 2537

Tóth SZ, Lőrincz T, Szarka A (2018) Concentration does matter: The beneficial and potentially harmful effects of ascorbate in humans and plants. Antiox Redox Signal 29: 1516–1533

Tóth SZ, Nagy V, Puthur JT, Kovács L, Garab G (2011) The physiological role of ascorbate as photosystem II electron donor: protection against photoinactivation in heat-stressed leaves. Plant Physiol 156:382–92

Tóth SZ, Oukarroum A, Schansker G (2020) Probing the photosynthetic apparatus noninvasively in the laboratory of Reto Strasser in the countryside of Geneva between 2001 and 2009. Photosynthetica 58: 560–572

Tóth SZ, Puthur JT, Nagy V, Garab G (2009) Experimental evidence for ascorbate-dependent electron transport in leaves with inactive oxygen-evolving complexes. Plant Physiol 149: 1568–1578

Tóth SZ, Schansker G, Garab G, Strasser RJ (2007) Photosynthetic electron transport activity in heat-treated barley leaves: The role of internal alternative electron donors to photosystem II. Biochim Biophys Acta 1767: 295–305

Tunc-Ozdemir M, Miller G, Song L, Kim J, Sodek A, Koussevitzky S, Misra AN, Mittler R, Shintani D (2009) Thiamin confers enhanced tolerance to oxidative stress in Arabidopsis. Plant Physiol 151: 421–432

Urano K, Yoshiba Y, Nanjo T, Igarashi Y, Seki M, Sekiguchi F, Yamaguchi-Shinozaki K, Shinozaki K (2003) Characterization of Arabidopsisgenes involved in biosynthesis of polyamines in abiotic stress responses and developmental stages. Plant Cell Environ 26: 1917–1926

Vardakou M, Salmon M, Faraldos JA, O’Maille PE (2014) Comparative analysis and validation of the malachite green assay for the high throughput biochemical characterization of terpene synthases. MethodsX 1: 187–196.

Vidal-Meireles A, Tóth D, Kovács L, Neupert J, Tóth SZ (2020) Ascorbate deficiency does not limit non-photochemical quenching in *Chlamydomonas reinhardtii*. Plant Physiol 182: 597–611

Volkening JD, Stecker KE, Sussman MR (2019) Proteome-wide analysis of protein thermal stability in the model higher plant *Arabidopsis thaliana*. Mol Cell Proteomics 18: 308–319

Vuckovic D (2012) Current trends and challenges in sample preparation for global metabolomics using liquid chromatography-mass spectrometry. Anal Bioanal Chem 403: 1523–154

Waditee-Sirisattha R, Shibato J, Rakwal R, Sirisattha S, Hattori A, Nakano T, Takabe T, Tsujimoto M (2011) The Arabidopsis aminopeptidase LAP2 regulates plant growth, leaf longevity and stress response. New Phytol 191: 958–969

Wang M, Jia Y, Xu Z and Xia Z (2016) Impairment of sulfite reductase decreases oxidative stress tolerance in *Arabidopsis thaliana*. Front Plant Sci 7:1843

Wang Z, Xiao Y, Chen W, Tang K, and Zhang L (2010) Increased vitamin C content accompanied by an enhanced recycling pathway confers oxidative stress tolerance in Arabidopsis. J Integr Plant Biol 52: 400–409

Warnes MGR, Bolker B, Bonebakker L, Gentleman R, Huber W (2016) Package ‘gplots’. Various R programming tools for plotting data. https://cran.r-project.org/package=gplots

Wheeler GL, Jones MA, Smirnoff N (1998) The biosynthetic pathway of vitamin C in higher plants. Nature 393: 365–369

Wickham H, Averick M, Bryan J, Chang W, McGowan LDA, François R, Grolemund G, Hayes A, Henry L, Hester J, Kuhn M, Pedersen LT, Miller E, Milton Bache S, Müller K, Ooms J, Robinson D, Paige Seidel D, Spinu V, Takahashi K, Vaughan D, Wilke C, Woo K, Yutani, H. (2019) Welcome to the Tidyverse. J Open Source Softw 4: 1686

Wickham H, Chang W, Wickham MH (2016) Package ‘ggplot2’. Create elegant data visualisations using the grammar of graphics. Version 2(1), 1–189. https://ggplot2.tidyverse.org/

Winter G, Todd CD, Trovato M, Forlani G, Funck D (2015) Physiological implications of arginine metabolism in plants. Front Plant Sci. 6:534

Wolucka BA, Van Montagu M (2003) GDP-mannose 3’,5’-epimerase forms GDP-L-gulose, a putative intermediate for the de novo biosynthesis of vitamin C in plants. J Biol Chem 278: 47483–47490

Xiao M, Li Z, Zhu L, Wang J, Zhang B, Zheng F, Zhao B, Zhang H, Wang Y, Zhang Z (2021) The multiple roles of ascorbate in the abiotic stress response of plants: Antioxidant, cofactor, and regulator. Front Plant Sci 12: 598173

Xu B, Sai N, Gilliham M (2021) The emerging role of GABA as a transport regulator and physiological signal. Plant Physiol 187: 2005–2016

Yabuta Y, Mieda T, Rapolu M, Nakamura A, Motoki T, Maruta T, Yoshimura K, Ishikawa T, Shigeoka S (2007) Light regulation of ascorbate biosynthesis is dependent on the photosynthetic electron transport chain but independent of sugars in Arabidopsis. J Exp Bot 58: 2661–2671

Yamamoto Y (2016) Quality control of photosystem II: The mechanisms for avoidance and tolerance of light and heat stresses are closely linked to membrane fluidity of the thylakoids. Front Plant Sci 7: 1136

Yerkes CT, Babcock GT (1980) Photosystem II oxidation of charged electron donors: surface charge effects. Biochim Biophys Acta 590: 360–372

Yin L, Fristedt R, Herdean A, Solymosi K, Bertrand M, Andersson MX, Mamedov F, Vener AV, Schoefs B, Spetea C (2012) Photosystem II function and dynamics in three widely used *Arabidopsis thaliana* accessions. PLOS One 7: e46206

Yoneyama T, Suzuki A (2020) Light-independent nitrogen assimilation in plant leaves: Nitrate incorporation into glutamine, glutamate, aspartate, and asparagine traced by 15N. Plants 9: 1303

Yu A, Xie Y, Pan X, Zhang H, Cao P, Su X, Chang W, Li M (2020) Photosynthetic phosphoribulokinase structures: Enzymatic mechanisms and the redox regulation of the Calvin-Benson-Bassham cycle. Plant Cell 32: 1556–1573

Yu Y, Wang J, Li S, Kakan X, Zhou Y, Miao Y, Wang F, Qin H, Huang R (2019) Ascorbic acid integrates the antagonistic modulation of ethylene and abscisic acid in the accumulation of reactive oxygen species. Plant Physiol 179: 1861–1875

Zechmann B (2018) Compartment-specific importance of ascorbate during environmental stress in plants. Antioxid Redox Signal 29: 1488–1501

Zechmann B, Stumpe M, Mauch F (2011) Immunocytochemical determination of the subcellular distribution of ascorbate in plants. Planta 233: 1–12

Zhang H, Whitelegge JP, Cramer WA (2001) Ferredoxin:NADP Oxidoreductase is a subunit of the chloroplast cytochrome b_6_f complex. J Biol Chem 276: 38159–38165

Zheng Q, Cheng ZZ, Yang ZM (2013) HISN3 mediates adaptive response of *Chlamydomonas reinhardtii* to excess nickel. Plant Cell Physiol 54: 1951–1962

